# Temporal Deconvolution of Mesoscale Recordings

**DOI:** 10.1101/2025.08.13.670164

**Authors:** Merav Stern, Eric Shea-Brown

## Abstract

Mesoscale calcium recordings, such as wide-field imaging, enable high-temporal-resolution recordings of neuronal activity across extensive brain regions. However, because these recordings capture light emitted by calcium indicator fluorescence, the underlying neural activity is obscured by the temporal dynamics of the indicators. Here, we develop and evaluate four deconvolution methods to recover neuronal spiking rates from fluorescence traces recorded in wide-field imaging. The methods—Dynamically-Binning (using adaptive discrete-time bins), Continuously-Varying (estimating smooth spiking rates), First-Differences (providing efficient estimation), and a modified Wiener Filter (robust to large fluorescence magnitude variations)-consistently outperform the previously used adapted Lucy-Richardson algorithm on both synthetic data and simultaneous fluorescence-spike recordings. Critically, we demonstrate that using raw fluorescence signals in advanced analysis methods can yield distorted results, including spuriously inflated correlations between brain regions’ activity. This highlights the necessity of accurate temporal deconvolution for reliable interpretation of brain activity.

## 2 Introduction

Recent developments in optogenetics allow the recording of high-temporal-resolution images of neuronal activity from the entire dorsal surface of the cortex in behaving animals (Makino et al. 2017, Chen et al. 2017, Allen et al. 2017), through the use of fluorescent calcium indicator molecules (Chen et al. 2013). Because these recordings capture large cortical areas at high temporal resolution, they are limited in their spatial resolution. Consequently, each pixel in the fluorescence images represents the combined activity of many thousands of neurons. The images can be collected continuously for hours at acquisition rates of dozens of hertz. Aggregating the images over time produces a fluorescence trace for each pixel. Inferring the time-varying neuronal activity from a given fluorescence trace is a challenging deconvolution problem.

Wide-field recording techniques date back several decades. They use a single-photon microscope and a camera to record changes in illumination associated with brain activity. Before optogenetic manipulations were available, wide-field recordings captured changes in luminance driven by blood flow changes in the brain (hemodynamic) (Masino et al. 1993, for example). Because wide-field recordings cover large fields of view, the signal captured in each pixel originates from multiple sources (blood vessels and many neurons). The resulting signal is thus sufficiently strong to be detected through a relatively thin skull, as is the case for mice, thereby avoiding invasive procedures and allowing recordings of *in vivo* brain activity in behaving animals. It is also possible to conduct wide-field recordings in anesthetized animals (Kalchenko et al. 2014) or *in vitro* slices.

In recent years, advances in optogenetics have further facilitated prolonged (multiple-day) neuronal recordings *in vivo* by integrating optogenetic manipulations with chronically implanted windows that provide access to a wide field of view. As a result, the wide-field imaging method has reemerged as a powerful tool for simultaneously capturing neural activity across multiple brain regions over prolonged periods. Consequently, wide-field imaging is now an established, commonly used standard technique to record neural activity (Silasi et al. 2016, Clancy et al. 2019, Mann et al. 2017, Musall et al. 2018, Chen et al. 2017, Allen et al. 2017, Aimon et al. 2015, Wekselblatt et al. 2016, Makino et al. 2017).

Despite these advancements, suitable methodologies for temporal deconvolution that infer neural spiking rate from fluorescence captured in wide-field recordings remain significantly lacking. In this study, we develop a mathematical framework that phrases the temporal inference problem in wide-field imaging as a tractable model. We introduce four innovative solutions: two statistical algorithms, one estimation calculation, and one filtering adaptation. We study their performance using both synthetic and recorded data.

In contrast, the problem of inferring a fluorescence trace that originates from the neural activity of a single neuron (typically obtained by two-photon imaging) has recently been considered by several authors, including Jewell & Witten (2018), Friedrich et al. (2017), and Pnevmatikakis et al. (2016). These authors use an autoregression model, originally proposed in Vogelstein et al. (2009), which associates the observed fluorescence 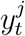 of a single neuron *j* at the *t*th timepoint with its unobserved calcium 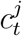 and emitted number of spikes 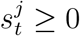,

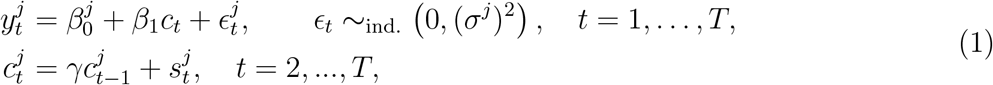

where *γ* ∈ (0, 1) is the rate of calcium decay. In words, (1) indicates that the calcium decays exponentially over time, unless there is a spike at the *t*th timepoint, in which case it increases; furthermore, the observed fluorescence is a noisy realization of the underlying calcium at each timepoint. In (1), 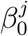 corresponds to the baseline fluorescence, which must be estimated; however, we can set *β*_1_ = 1 without loss of generality, as this simply amounts to scaling the fluorescence by a constant factor.

To fit the model (1), Friedrich et al. (2017), Jewell & Witten (2018), and Jewell et al. (2019) solve the optimization problem

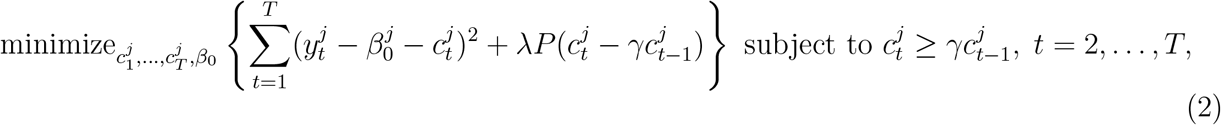

where *λ* is a nonnegative tuning parameter, and *P* (·) is a penalty function designed to induce sparsity by encouraging 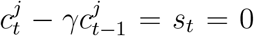 so that at most timepoints no spike is estimated to occur. Friedrich et al. (2017) makes use of an *ℓ*_1_ penalty (Tibshirani 1996), while Jewell & Witten (2018) and Jewell et al. (2019) instead use an *ℓ*_0_ penalty.

In mesoscale recordings, such as wide-field imaging, fluorescence traces typically reflect the activity of a large population of neurons rather than a single neuron. Consequently, single-neuron deconvolution solutions, which assume that there are no spikes at most timepoints, cannot be applied directly. In this manuscript, we develop an extension of the model (1) and its corresponding optimization problem (2) to the setting of wide-field calcium imaging recordings. We propose novel solutions that fit our innovative approach for deconvolving large-scale neural activity. We apply our new approach for the deconvolution of wide-field calcium imaging recordings to data from V1 (Clancy et al. 2019) and the whole dorsal surface (Musall et al. 2018, Gilad et al. 2018).

## 3 Results

### 3.1 Novel Temporal Deconvolution Methods

We present and evaluate four deconvolution methods for inferring the spiking rate *r* from a fluorescence trace 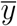 obtained by mesoscale recordings, such as wide-field imaging. All our methods adhere to the model we develop in the following section for fluorescence resulting from aggregated neural activity, Equation (3). In our first approach, we formulate an optimization problem that corresponds to our model. Solutions to this problem yield two deconvolution methods: the ‘Dynamically-Binning’ algorithm and the ‘Continuously-Varying’ algorithm. Our second approach infers the spiking rate point-by-point in time according to our model and then estimates (and removes) the associated noise from the overall result. The solution obtained by this approach is the ‘First-Differences’ deconvolution method. In our third approach, we adapt a known filter, the ‘Wiener Filter’. Our model is used in this approach to help define the filter, which removes the noise in the frequency domain, allowing us to restore the spiking rate in the temporal domain.

After presenting our novel approaches, we compare their performance with that of the ‘Lucy-Richardson’ algorithm, which has previously been employed to deconvolve fluorescence captured in wide-field recordings (Wekselblatt et al. 2016). ^1^

### 3.2 A Model for Mesoscale Imaging of Neuronal Activity

We extend here the model (1) to wide-field recordings. To begin, we consider the model (1) for multiple *j* = 1, …, *p* neurons simultaneously. In this setting, the observed fluorescence trace is the sum of the fluorescence associated with each of *p* neurons recorded at a given time in a given pixel. Summing (1) across the *p* neurons yields the model

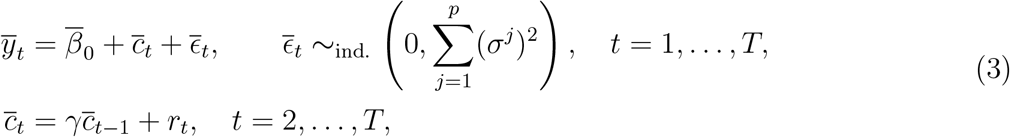

where 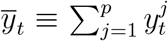 is the total observed fluorescence of the *p* neurons at the *t*th timepoint, and where 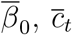, and 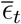 are the total baseline fluorescence, total calcium, and total noise at the *t*th timepoint, respectively.

In (3), 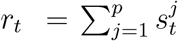 represents the aggregate spiking activity of all neurons contributing to the fluorescence at the *t*th timepoint. We refer to *r*_*t*_ as the *spiking rate*. It captures the total spike-driven increase in calcium across this neural population. This notation emphasizes the key shift from single-neuron spike inference, where *s*_*t*_ represents sparse spike events, to mesoscale population-rate inference, where *r*_*t*_ represents the aggregate activity of many neurons. Because *p* (the number of neurons contributing to the fluorescence) is potentially quite large, on the order of hundreds and thousands of neurons, we do not expect *r*_*t*_ to be sparse. Instead, we expect *r*_*t*_ to take on similar values at nearby timepoints, for the most part. Hence, we do not expect the spiking rate to vary over time in an arbitrary way. We will explore this point in greater detail in the next section.

The total baseline fluorescence, 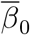, typically depends on the processing performed on the observed fluorescence (see, e.g. the Δ*F/F* preprocessing of Chen et al. (2017)).

In building the model (3), we implicitly assume that the calcium decay rate, *γ*, is fixed across neurons and over time. This is a constructive approximation, as it allows us to formulate an analytically tractable model and derive solvable deconvolution frameworks from it. However, it is possible for calcium decay rates to exhibit slight variations across neurons and within the same neuron over time.

For example, Chen et al. (2013) reported deviations of up to 10% between neurons of the same type and within the same neuron over time, depending on the neuron’s activity. Nevertheless, methods for single-neuron spike inference (Jewell & Witten 2018, Friedrich et al. 2017, Pnevmatikakis et al. 2016) assume a constant decay rate over time. These methods have been shown to provide reliable estimates of neural activity and are widely used. In the wide-field setting, we make a similar assumption across neurons. Consequently, the aggregated calcium concentration *c* inherits its effective decay rate from the single-neuron decay rate. Our analysis of parallel fluorescence and spike counts recordings in Section 3.13 demonstrates that indeed, the decay rates minimizing inference of mesoscale activity error fall within the experimentally observed range for single neurons. Although a fixed *γ* does not capture all biological variability, we show that this approximation performs well and is empirically supported for modeling and deconvolution. We discuss this point further when we analyze the parallel recordings in Section 3.13.

### 3.3 Optimization Problem

The model (3) leads naturally to the optimization problem

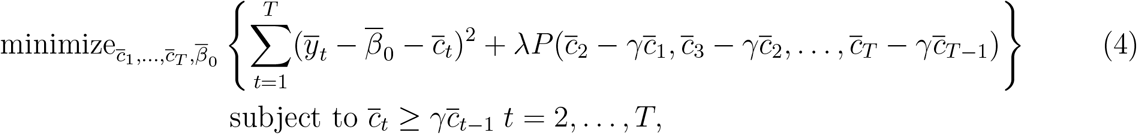

which closely resembles the optimization problem used to deconvolve the fluorescence trace for a single neuron (Friedrich et al. 2017, Jewell & Witten 2018, Jewell et al. 2019). For that task, the authors considered the use of a sparsity-inducing penalty because a neuron is not expected to spike at most timepoints. In contrast, here we are considering a model in which 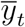 represents the total fluorescence at the *t*th timepoint summed over *p* neurons and in which 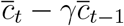 represents the spiking rate *r*_*t*_ at the *t*th timepoint across all *p* neurons. Therefore, in the context of wide-field imaging, we do not want *P* (·) to be a sparsity-inducing penalty. Instead, we want *P* (·) to encourage adjacent timepoints to, for the most part, have similar values of *r*_*t*_. A penalty *P* (·) that encourages spiking rate solutions *r*_*t*_ to maintain similar values at adjacent timepoints can take the form:

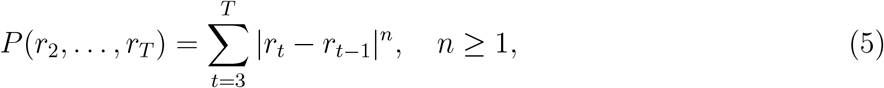

To incorporate penalty (5) into the optimization problem (4), we reparameterize the latter in terms of *r*_1_, …, *r*_*T*_. We use 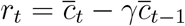 for *t* = 2, …, *T* as defined in (3), and 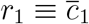 which is strictly a definition intended for notational convenience. This reparameterization can be expressed in matrix form as 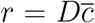, where *D* is a *T × T* full-rank matrix with 1 on the diagonal and −*γ*’s just below the diagonal. This leads to a rephrasing of the optimization problem (4) as follows:

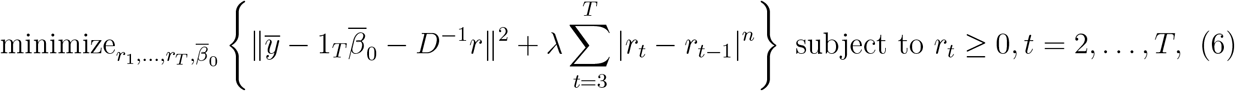

where 1_*T*_ is a vector of length *T* with all elements equal to 1.

We observe that a solution 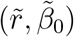 to the optimization problem (6) obeys:

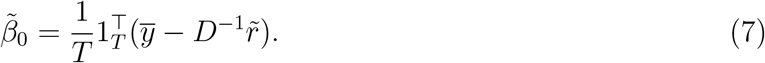

By substituting it into the optimization problem (6), we can obtain the following equivalent optimization problem for the spiking rate:

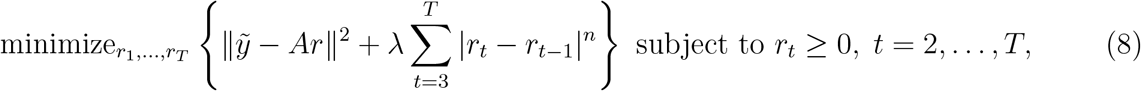

where 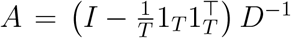 and 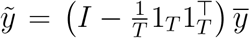, with *I* is the *T × T* identity matrix and 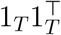 is a *T × T* matrix of ones.

Note that 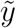 is the mean-centered version of 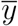, and *A* is the column-mean-centered version of *D*^−1^, which relates the calcium to the spiking rate via 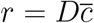. Optimization problem (8) is simpler to solve since it depends solely on the spiking rate. Thus, we can first obtain a solution to the spiking rate via solving (8) and then compute the intercept 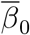 using its closed-form expression in (7). In Sup. information S1.1 and S1.2, we prove that solving the optimization problem (8) and calculating 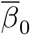 via (7) is equivalent to solving the optimization problem (4) with the penalty (5) that was chosen to fit wide-field imaging properties.

### 3.4 The Dynamically-Binning Deconvolution Method

We consider the penalty (5) with *n* = 1.

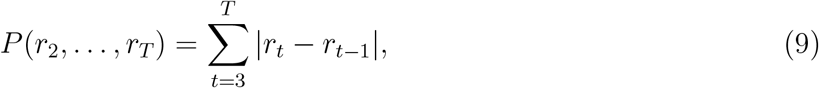

which is a *fused lasso*, or *total variation denoising*, penalty (Tibshirani et al. 2005, Tibshirani et al. 2012, Condat 2013). This penalty limits frequent abrupt changes in the spiking rate. In addition, and unique to this penalty, it encourages the spiking rate, *r*_*t*_, to be constant over time, with only occasional changepoints, as shown in Figure 1(a)i. This yields estimates of the spiking rate that are constant within a bin, where the bins themselves are adaptively estimated from the data. With penalty (9) and as discussed in Section 3.3, we can find the dynamically-binned spiking rate from a fluorescence trace recorded using wide-field imaging by solving

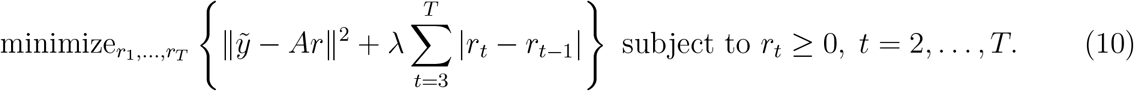

**Figure 1:**
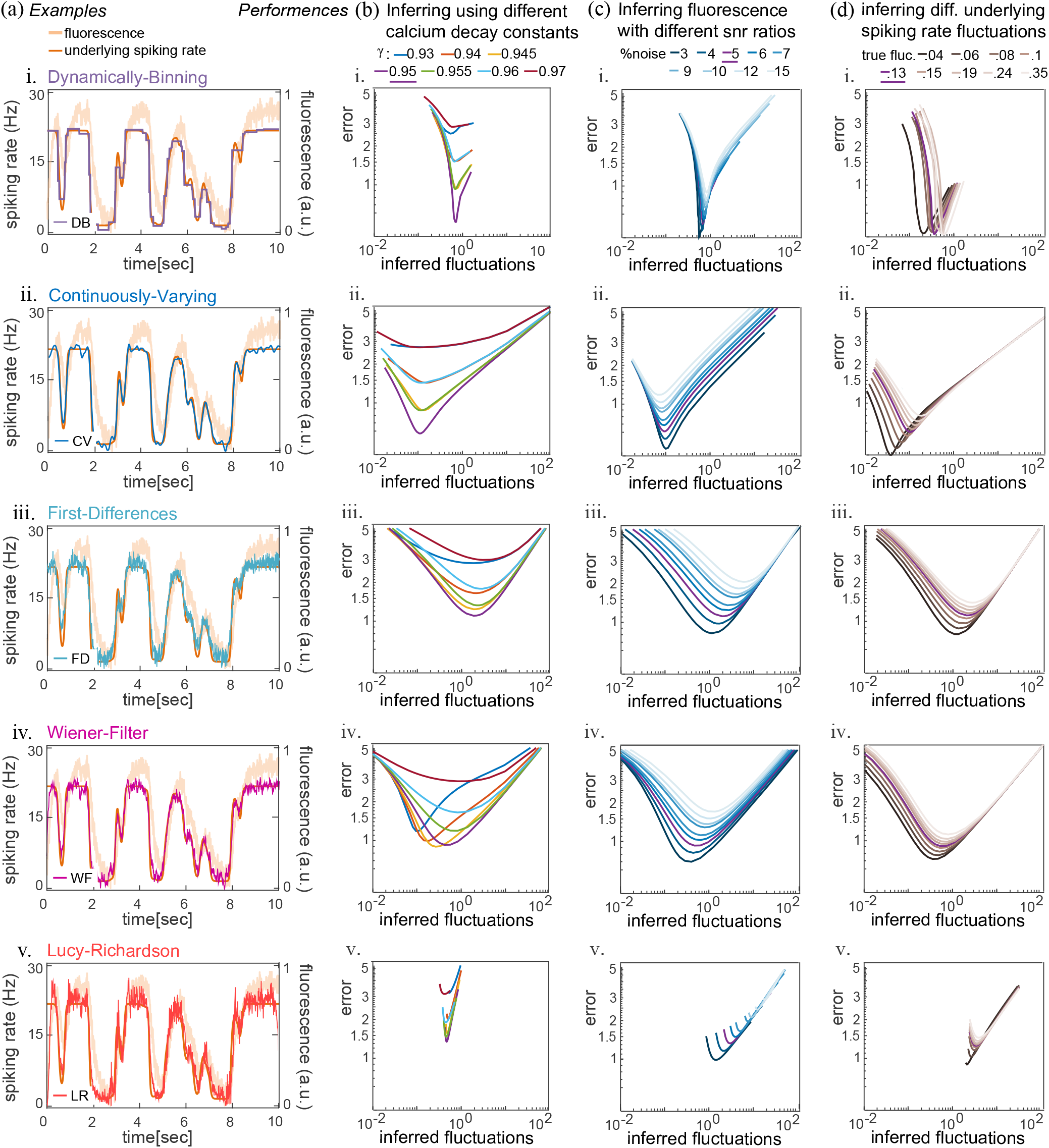
Detailed Simulated Data Results. Caption of Figure 1 (a) An example of an underlying simulated spiking rate trace (orange line, see Supplementary Information S2 for simulation details) and its resulting fluorescence trace (ivory line, calculated using the model (3)). The inferred spiking rates by the i. Dynamically-Binning, Algorithm 1, ii. Continuously-Varying, Algorithm 2, iii. First-Differences, Equation (15), iv. Wiener-Filter, Equation (17), and v. Lucy-Richardson deconvolution, appear in purple, blue, light blue, magenta, and red lines, respectively. (b). The mean error, Equation (18), between the underlying and the inferred spiking rate by each of the methods: i. DB, ii. CV, iii. FD, iv. WF, and v. LR. Each line corresponds to deconvolution using a different calcium decay *γ*. The differences in fluctuations (19) are due to using different parameter values (penalty *λ* in DB and CV, smoothing in FD or inverse SNR in WF) with smaller values of this parameter leading to higher fluctuations and vice versa. In LR, smaller parameter values (number of iterations) decrease fluctuations and vice versa. All methods are consistent in the sense that the best inference (with minimum error) is achieved with *γ* = 0.95 (purple line), which was used to simulate the data. (c-d) The mean error, Equation (18), between the underlying and the inferred spiking rate as a function of the mean inferred rate fluctuations, Equation (19). (c) Each line corresponds to a different amount of noise *σ* present in the simulated fluorescence traces relative to the calcium signal *c* in it, 0.03 ≤ *σ/*| max*c* − min *c*| ≤ 0.15. (d) Each line corresponds to a different underlying spiking rates dataset with a different fluctuation level. (c-d) For the deconvolution *γ* = 0.95 was held fixed. Increasing *λ*, for i. DB and ii. CV, the number of smoothing points, for iii. FD, inverse SNR, for iv. WF, or limiting the iteration number, for v. LR, decreased the inferred fluctuations and vice versa. Across almost all conditions, DB performs the best, including its ability to maintain fixed inferred fluctuations despite increased noise, as demonstrated in (c)i. LR performs the poorest, and FD the second-poorest, with larger inferred fluctuations (in addition to larger errors) compared to the other methods across conditions. This manifests in a relatively noisy inferred spiking rate (as evident in (a)v. red line and (a)iii. light blue line, respectively). A quantified comparison between the quality of the methods’ performances on synthetic data appears in Figure 2.

This problem is a convex optimization problem (Boyd & Vandenberghe 2004) that can be efficiently solved for the global optimum. Since its penalty is non-differentiable, we utilize the proximal gradient descent algorithm for solving it, which results in Algorithm 1. The details of finding Algorithm 1 are further explained in Sup. information S1.3.

#### Algorithm 1

Dynamically-Binned Rate Deconvolution: Solving (10)

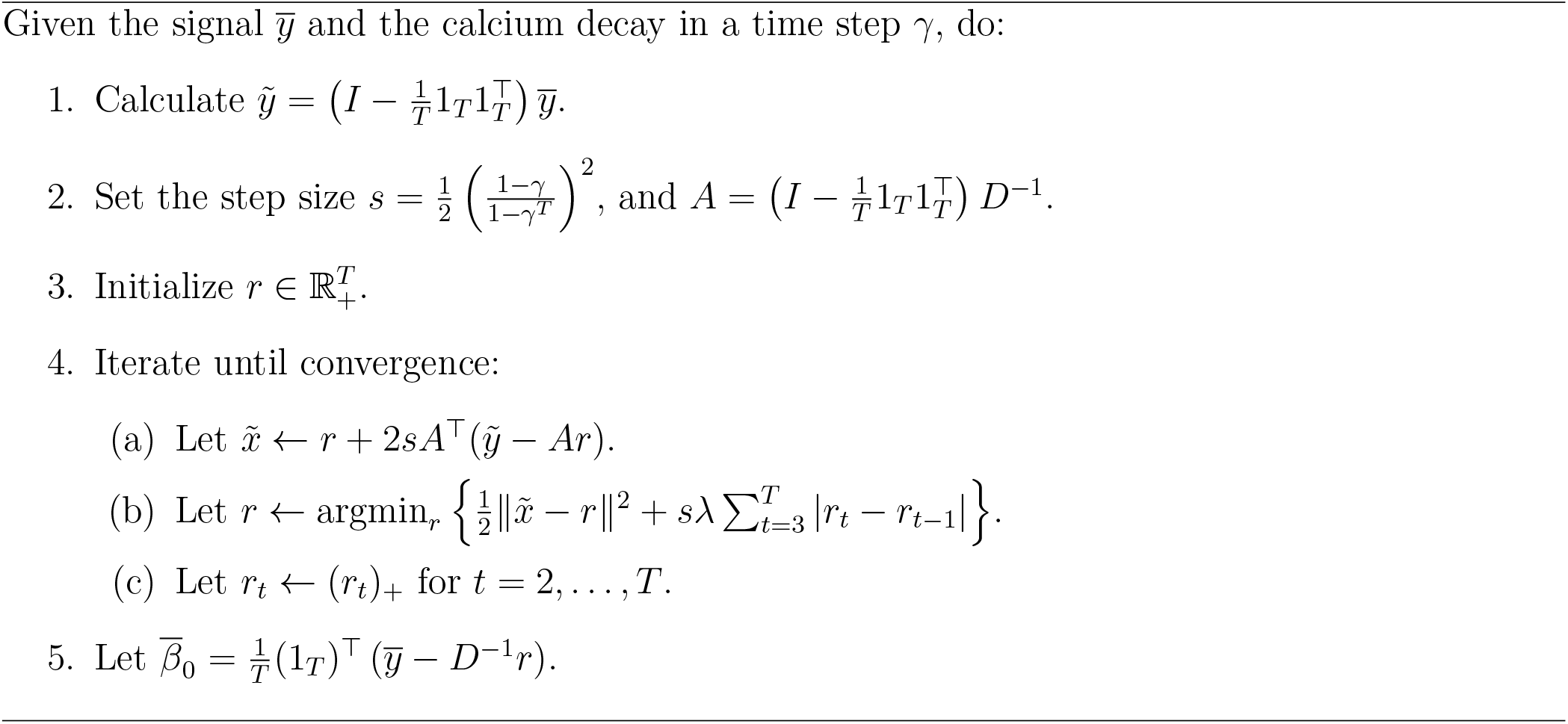

### 3.5 The Continuously-Varying Deconvolution Method

We consider the penalty (5) with *n* = 2

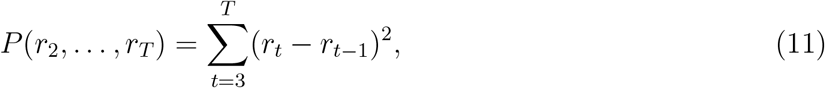

This penalty restricts sudden shifts in the spiking rate. It does so by encouraging the spiking rate to continuously vary over time, as shown in Figure 1(a)ii.

It is convenient to express the penalty (11) in matrix form as

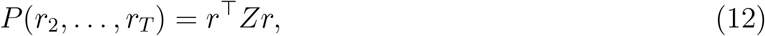

where *Z* is defined by

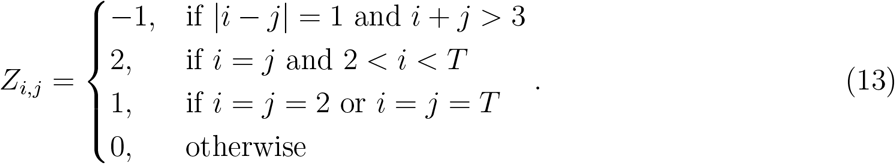

With the penalty (12) and as discussed in Section 3.3, we can find the continuously-varying spiking rate from a fluorescence trace recorded using wide-field imaging by solving

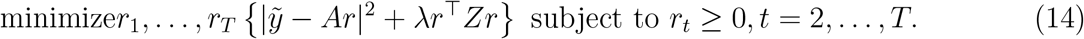

This problem is a convex optimization problem (Boyd & Vandenberghe 2004) with a differentiable penalty and non-negativity constraints on its solutions. To solve it, we again use the proximal gradient descent algorithm^2^. This results in Algorithm 2. The details of finding Algorithm 2 are further explained in Sup. information S1.6.

#### Algorithm 2

Continuously-Varying Rate Deconvolution: Solving (14)

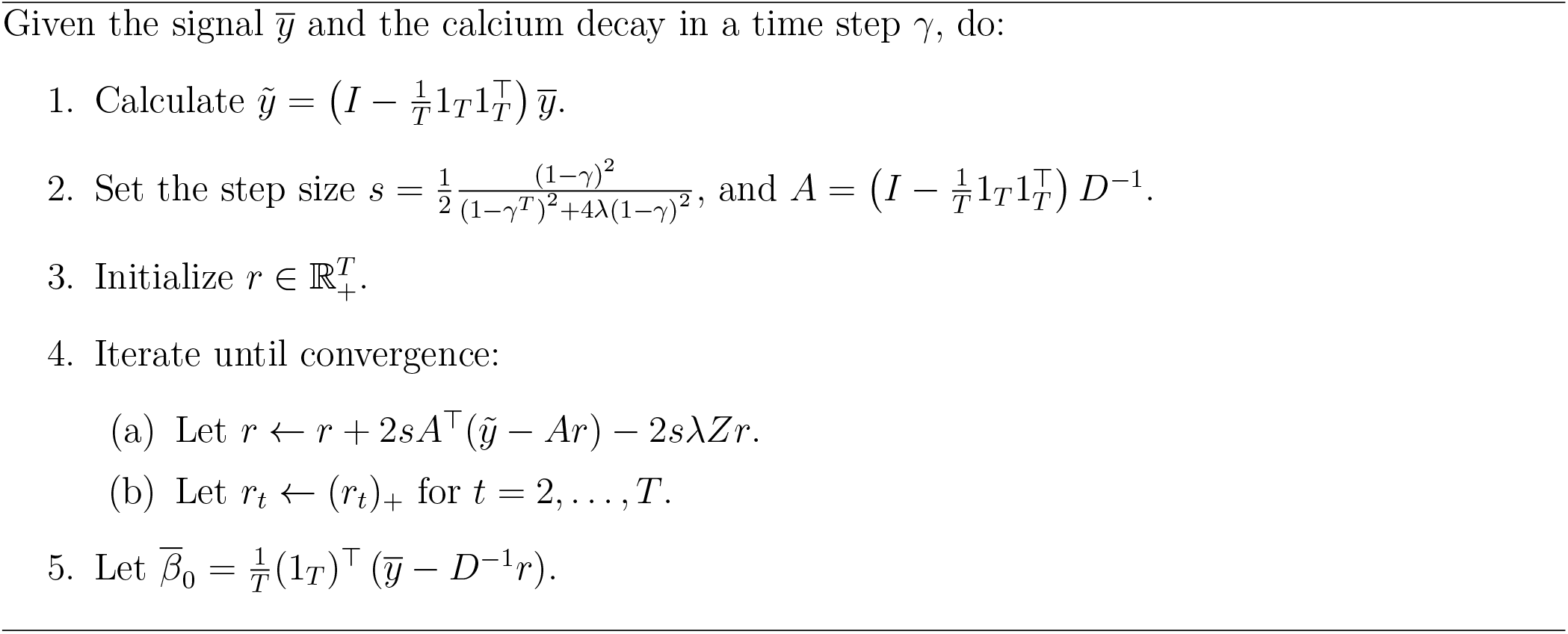

### 3.6 The First-Differences Deconvolution Method

This method assumes that the calcium concentration is directly represented by the fluorescence. Hence, the spiking rate can be found by assuming 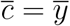 in (3), which yields

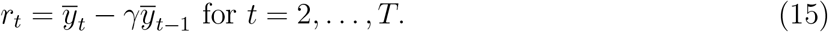

In the special case of *γ* = 1, this method is equivalent to estimating the spiking rate by computing the first differences (derivative) of the observed fluorescence.

The spiking rate obtained from (15) can be noisy and may not satisfy the non-negativity constraint. To address these issues, we add two additional steps to the First-Differences method. First, a moving average window is applied to smooth the estimated spiking rate following (15). Second, a mean shift is introduced to prevent negative rates.

Overall, this method is a natural, simple alternative to our minimization-based approach. It bypasses the need to estimate the underlying calcium concentration and replaces the penalties in (9) and (11) with a simple smoothing *post hoc* procedure. An example of a spiking rate obtained using this method is shown in Figure 1(a)iii.

### 3.7 The Wiener-Filter

This method utilizes the Wiener filter (Wiener 1964), a standard signal processing technique, to infer spiking rates from mesoscale fluorescence recordings. The Wiener filter provides an effective approach to mitigate noise within a filtered signal (Dogariu et al. 2021), operating under the assumption that the ratio of noise variance to signal variance is fixed and known. Since this ratio is not known a priori for mesoscale fluorescence recordings, we treat it as a tuning parameter, which we denote as *k*.

To utilize the Wiener filter, an explicit formulation of the filter is required. According to our model (3), the filter *h*(*t*) of the spiking rates by the calcium dynamics is given by:

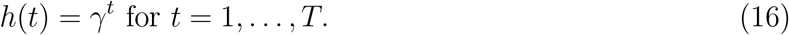

Following the Wiener filter technique, we determine the spiking rate in the frequency domain as follows:

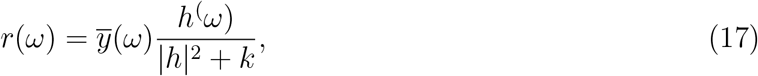

where 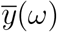 is the Fourier transform of 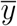, *k* is the tuning parameter representing the unknown ration of noise variance to signal variance, *h*(*ω*) is the conjugate of the filter (16) Fourier transform and 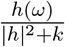 is the Wiener filter itself. The spiking rate *r*(*t*) is then found by the inverse Fourier transform of the result *r*(*ω*) in (17). An Example of a spiking rate inferred by the Wiener-Filter is shown in Figure 1(a)iv.

### 3.8 Comparison to the Lucy-Richardson Algorithm

The Lucy-Richardson deconvolution method is an iterative algorithm that was originally developed in astrophysics to restore light sources from filtered, blurred images (Lucy 1974, Richardson 1972). This method assumes that the blurring is due to a convolution filter *f* and additive noise. It operates by maximizing the likelihood of the original and denoised images, based on the assumption that the noise follows a Poisson distribution. In the context of wide-field calcium imaging, the Lucy-Richardson algorithm has been adapted for temporal deconvolution using a one-dimensional filter that describes the calcium decay process, defined as *h*_*t*_ = *γ*^*t*^ (Equation (16)) (Wekselblatt et al. 2016). An Example of a spiking rate inferred by the Lucy-Richardson algorithm is shown in Figure 1(a)v.

The performance of the Lucy-Richardson algorithm may depend on the number of iterations performed (resembling the role of the penalty *λ* in Dynamically-Binning and Continuously-Varying, the number of smoothing points in First-Differences, or the inverse SNR *k* parameter in Wiener-Filter). We varied this number in Figure 1(b-d)v to facilitate easy visual comparison between the Lucy-Richardson algorithm and all of our other proposed deconvolution methods described above. We discuss the results in Section 3.11 in detail.

### 3.9 Performance Measures

To quantitatively assess the performance of the different deconvolution methods, we can calculate the difference between the inferred spiking rate by a deconvolution method *r*^*est*^ and the underlying spiking rate *r* (when the latter is known). This gives an error measure:

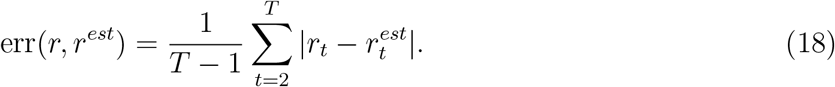

Note that since a solution to (6) is unique only up to a constant shift (see Sup. information S1.8), we compute (18) after subtracting the mean from both *r*_*t*_ and 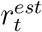 for *t* = 2, …, *T*.

A second useful measure is the level of fluctuations, which indicates how rapidly a trace *x* changes. It can be naturally defined by

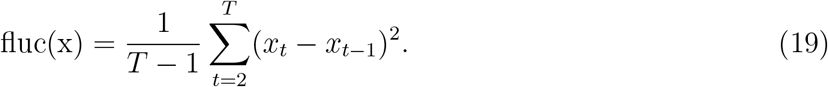

where *x* represents the fluorescence *y*, calcium *c* or spiking rate *r* of either the underlying or the inferred solution. Note that when *x* represents the spiking rate, Equation (19) is equivalent (up to a normalization factor) to the penalty term used in problem (6) with *n* = 2 (Equation (11)).

### 3.10 Method Notation

In the following sections, we provide a detailed comparison of the performance of the deconvolution methods across different conditions and datasets. To ease readability, we use the following short-hand method names in the main text: Dynamically-Binning as Dynbin, Continuously-Varying as Convar, First-Differences as Firdif, Wiener Filter as Wiener, and Lucy-Richardson as Lucy. In figures and captions, where space is limited, we employ the more compact abbreviations DB, CV, FD, WF, and LR, respectively.

### 3.11 Results on Simulated Data

We tested the performance of the deconvolution methods on simulation data. Our data includes 10,000 underlying spiking rate traces recorded at 20 Hz, generated by an artificial neural network we built (details of the network are given in Sup. information S2). From the simulated spiking rates, we calculated their corresponding calcium 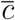 and fluorescence 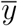 traces, using the model (3). We used a <u>c</u>alcium decay constant of *γ* = 0.95 and a noise variance 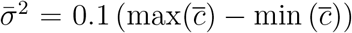. We choose 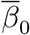 randomly.

We applied the deconvolution methods to the fluorescence traces we generated and compared the inferred spiking rates to the underlying spiking rates. Overall, all the methods yielded reasonable results, with some performing better than others. They all inferred spiking rates that resemble the underlying rates, but with varying degrees of precision (examples are shown in Figure 1(a)i-v).

Using different calcium decay rate values *γ* and parameter values (penalty weight *λ* for Dynbin and Convar, number of smoothing points for Firdif, inverse SNR *k* for Wiener, and number of iterations for Lucy) resulted in different average error (18) and average inferred spiking rate fluctuations (19) (shown in Figure 1(b)i-v, where different lines correspond to using different calcium decays for deconvolution). Higher penalty weights for Dynbin and Convar decreased inferred spiking fluctuations, and vice versa. More smoothing points for Firdif, higher inverse SNR for Wiener, or fewer iterations for Lucy also decreased inferred spiking fluctuations. All the methods were found consistent in the sense that they produce a minimum error when using the calcium decay value that matches the fluorescence simulations (Figure 1(b), purple lines). Inferred spiking rates obtained by Dynbin were the most accurate. Convar yielded similar results, with only slightly elevated errors. The inferred spiking rates by Lucy were the least accurate (shown in Figure 1 (b) i-v)

We extended the simulated data by incorporating various levels of noise (*ϵ* in the model (3)) into the fluorescence traces that were generated from otherwise the same spiking rate traces as before. We kept the calcium decay *γ* = 0.95 fixed during deconvolution and adjusted the parameter values (penalty weight *λ* for Dynbin and Convar, number of smoothing points for Firdif, inverse SNR *k* for Wiener, or number of iterations for Lucy). The increase in noise made deconvolution more challenging for all methods, as evidenced by the increase in error between the inferred and underlying spiking rates (see Figure 1(c)). Note that, Convar inferred spiking rates exhibited decreased fluctuations with increased noise, whereas Firdif, Wiener, and Lucy inferred spiking rates exhibited increased fluctuations with increased noise (this is despite the fact that the underlying spiking rates remained unaffected by the varying noise levels). Dynbin managed to maintain stable fluctuations in the inferred spiking rates across the different noise levels.

We also tested the deconvolution methods on fluorescence traces generated from spiking rates with varying levels of fluctuations. We controlled the underlying spiking rate fluctuations by adjusting the gain *g* in the neural networks we have built (see Sup. information S2). All methods showed increased inferred spiking rate fluctuations corresponding to the increase in underlying spiking rate fluctuations. Although Convar, Firdif, Wiener, and Lucy exhibited growing errors in the inferred spiking rates as fluctuations increased, Dynbin managed to maintain the lowest error consistently across all fluctuation levels (Figure 1(d)).

Note that the performance of the Lucy-Richardson algorithm depends only weakly on the number of iterations. This is evident from the small variability within each curve in Figure 1(b-d)v. In contrast, the other methods show much larger variability across their corresponding parameters, the penalty *λ* in DynBin and ConVar, the number of smoothing points in FirDif, and the inverse SNR parameter *k* in Wiener, as evident from the large variability within each curve in Figure 1(b-d)i-iv.

Consistent with this weak dependence, we found that executing Lucy-Richardson with 10 iterations yields the best results (with minimal error) across all consecutive datasets and conditions. In rare cases, 8, 9, 11, or 12 iterations yielded minimal error, but the difference compared to 10 iterations was negligible. This indicates that the optimal number of iterations for the Lucy-Richardson algorithm is not strongly related to the nature of the data, but possibly an intrinsic property of the algorithm or its interaction with the temporal calcium decay filter; it is worth noting that 10 iterations is also the default setting for the Lucy-Richardson algorithm in different programming packages. Therefore, for later analysis, we set the number of iterations to 10. This implies that there is no parameter to tune when using the Lucy-Richardson algorithm for wide-field imaging inference. This can be a significant advantage of this algorithm as it shortens and simplifies its implementation. However, as we demonstrate below (and regardless of whether we chose or what we chose as the iteration number), its performance is worse than all of our other suggested deconvolution methods across almost all datasets, conditions, and measures.

### 3.12 Comparison of Simulated Data Results

To better understand the results above, we compared the methods’ best performances across the different conditions (see Figure 2).

**Figure 2:**
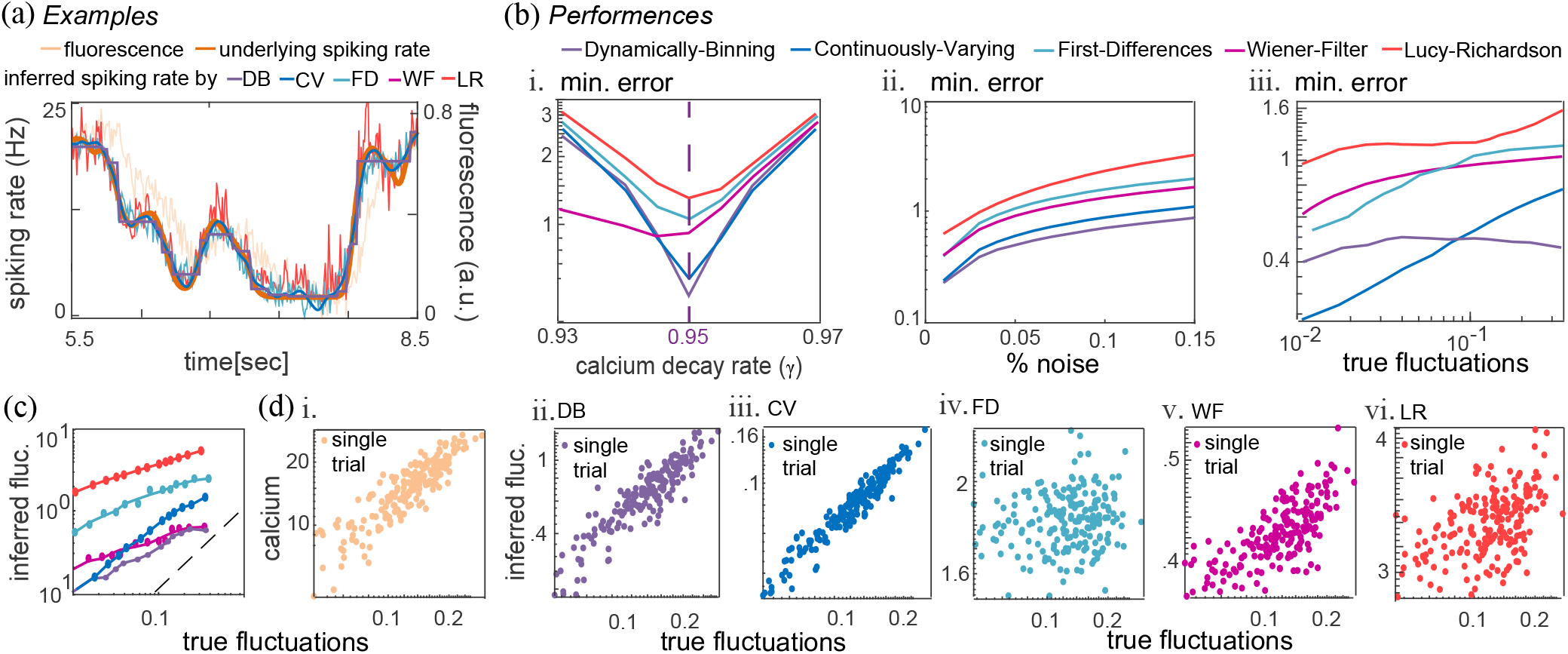
Comparison of Simulated Data Results.(a) An example of an underlying simulated spiking rate trace (orange line, see Sup. information S2 for simulation details) and its resulting fluorescence trace (ivory line, calculated using the model (3)). The inferred spiking rates by the Dynamically-Binning algorithm (purple line), Continuously-Varying algorithm (blue line), First-Differences method (light blue line), Wiener-Filter (magenta), and the Lucy-Richardson deconvolution (red line) appear on top. (b) The minimal mean error between the underlying and the inferred spiking rate for each method: DB purple line, CV blue line, FD light blue line, WF magenta line, LR red line, as a function of: i. deconvolution with different calcium decay values, ii. deconvolution of fluorescence simulated with different ratios between the added noise amplitude (measured by the standard deviation of the noise) and the calcium signal amplitude (measured by the range of the calcium absolute values), iii. deconvolution of fluorescence generated from simulated underlying spiking rates with varying fluctuations. The minimal error was determined by finding the best parameter values (penalty *λ* for BD and CV, smoothing for FD, inverse SNR for WF, and iteration number for LR) that yield the minimum error for each method in otherwise equal conditions. In other words, the minima point in each of the lines in Figures 1(b-d) construct plots (b)i-iii. (c) The mean inferred spiking rate fluctuations as a function of the mean underlying spiking rate fluctuations. The black dashed line marks *x* = *y*. (d) As a function of the true (underlying) spiking rate fluctuations of 200 example traces (a dot represents each trace’s fluctuations), the following appear: i. The fluctuation level of the calculated calcium trace from the underlying spiking rate (using the model (3)); ii-vi. The fluctuation level of the inferred spiking rate by ii. DB, iii. CV, iv. FD, v. WF, vi. LR.

The Lucy algorithm produced the noisiest and most fluctuating solutions (an example is shown in Figure 2(a) red line). Firdif method yielded the second noisiest fluctuating solutions (an example is shown in Figure 2(a) light blue line). This is also evident in the highest and second-highest errors for Lucy and Firdif, respectively (Figure 2(b)i), and the highest and second-highest inferred fluctuations (depicted in Figure 2(c)).

We also examined which deconvolution method best preserves the (fluctuations) structure of the data. To do so, we calculated the correlation between the fluctuations, Equation (19), of all simulated underlying spiking rate traces and the fluctuations of their corresponding inferred spiking rate traces (for short, we refer to the latter as inferred fluctuations). We expect “well-behaved” deconvolution methods to infer spiking rates with fluctuations that match those of the fluorescence traces from which they were inferred (whether high, medium, or low). The calcium dynamics (together with the noise) constitute the fluorescence signal. It filters the neural dynamics but maintains information about it, including an indication of its fluctuation level. This is evident in Figure 2(d)i where the underlying spiking rate fluctuations and their corresponding calcium trace fluctuations are visibly highly correlated (with 0.88 Pearson correlation). Therefore, a deconvolution should preserve the fluctuation structure in the data, allowing us to infer high-(or low-) spiking-rate fluctuations from high-(or low-) fluctuating fluorescence. In our tests, Convar best preserved the (fluctuations) structure, with a correlation of 0.95 between the underlying and inferred spiking rate fluctuations (Figure 2(d)ii). Dynbin was the second best with a correlation of 0.89 (Figure 2(d)iii), Wiener was third with a correlation of 0.58 (Figure 2(d)iv),Lucy fourth with 0.37 (Figure 2(d)vi) and Firdif almost didn’t preserve the fluctuations structure at all with a correlation of 0.13 (Figure 2(d)v).

In Table 1, we summarize the performance of the deconvolution methods across all the criteria above and when inferring real data, which we discuss in what follows.

**Table 1:** Summary of deconvolution method performance across datasets and evaluation metrics. For each dataset and metric, the table assigns each method to one of four qualitative performance categories: Exc (Excellent), Good, Fair, or Poor. Runtime is summarized as Fast, Moderate, or Slow.

|  | Simulations |  | With parallel spike counts |  |  | Fluorescence only |  |  | Runtime |
| --- | --- | --- | --- | --- | --- | --- | --- | --- | --- |
| Method | Error<br>Fig. 2b | Fluc.<br>corr.<br>Fig. 2d | Error<br>Fig. 4a | Fluc.<br>corr.<br>Fig. 4c | Param.<br>consist.<br>Fig. 4b | Error<br>Fig. 5e,<br>6a | Fluc.<br>corr.<br>Fig. 5e,<br>6a | Remove<br>ghosts<br>Fig. 5a,<br>6a |  |
| Dynamical Binning | Exc. | Good | Good | Good | Fair | Fair | Poor | Poor | Slow |
| Continuously Varying | Exc. | Exc. | Good | Good | Exc. | Good | Exc. | Exc. | Slow<sup>3</sup> |
| First Differences | Fair | Poor | Fair | Fair | Poor | Good | Exc. | Fair | Fast |
| Wiener Filter | Fair | Fair | Exc. | Exc. | Good | Good | Good | Good | Moderate |
| Lucy-Richardson | Poor | Poor | Poor | Poor | Exc. | Poor | Poor | Fair | Moderate |

### 3.13 Parallel Fluorescence and Spike Count Recordings

Here, we compare the performance of the deconvolution methods on fluorescence traces with parallel recording of spike counts originating from the same location as the fluorescence. The recordings were collected by Kelly Clancy; see Clancy et al. (2019) Supplementary material Figure 4. They include 5 datasets relevant to our study, each comprising about an hour of parallel recordings, with a single spike-count trace and its corresponding (single) fluorescence trace. To study these datasets, we segmented their long traces into many shorter, equal segments of 20 seconds each. See a full description of the datasets and our prior-to-deconvolution analysis of them in Sup. information S3.

It is paramount to keep in mind that the spike counts recorded in Clancy et al. (2019), in parallel to the fluorescence, are an indication only of the true underlying spiking rate that drives the recorded fluorescence. Vast amounts of neurons contribute to a fluorescence trace in mesoscale recordings, such as wide-field imaging, beyond the ability of a silicon probe to capture (and no currently available technology can record the precise true underlying spiking activity contributing to the fluorescence in such imaging techniques).

Albeit the spike counts provide an estimate of the true underlying spiking rates, which is sufficient for our study. This is due to several reasons. First, neural activity tends to be highly correlated, particularly among neurons in close proximity. Since single recorded fluorescence traces in wide-field imaging originate from nearby neurons, their activities are generally highly correlated. Hence, spike counts from a subset of these neurons represent the larger surrounding population decently.

Moreover, wide-field imaging captures fluorescence originating from a large neural population, which primarily reflects correlated activity. The uncorrelated, unique activities of individual neurons generate a baseline activity in the cortex, which translates into baseline fluorescence (typically removed by performing a Δ*F/F*, subtracting *β*_0_, or through other preprocessing methods). This uncorrelated activity also manifests as a relatively stable baseline activity in spike counts, leaving the correlated activity to drive significant changes in spike counts, which describes well the population-level activity captured by the wide field.

Also note that the deconvolution process is independent of the spike count and relies solely on fluorescence data. This means that partially inaccurate spike counts do not affect the deconvolution results across any of our methods. The spike counts are only used to calculate the error between them and the inferred spiking rates. In a dataset where the spike count is unrelated to the fluorescence, the best deconvolution results (with minimum error) are obtained using either the largest or smallest penalty (or smoothing) available. This leads to a constant inferred spiking rate solution or excessive noise (not shown), respectively. In all our methods and across all datasets examined in this section, the optimal solution is achieved with a medium penalty value (as illustrated by the convex error curves in Figure 3(c)). This implies that the data is sufficient to convey a meaningful signal and that our deconvolution methods are effective enough in capturing information about the spike counts from the fluorescence.

**Figure 3:**
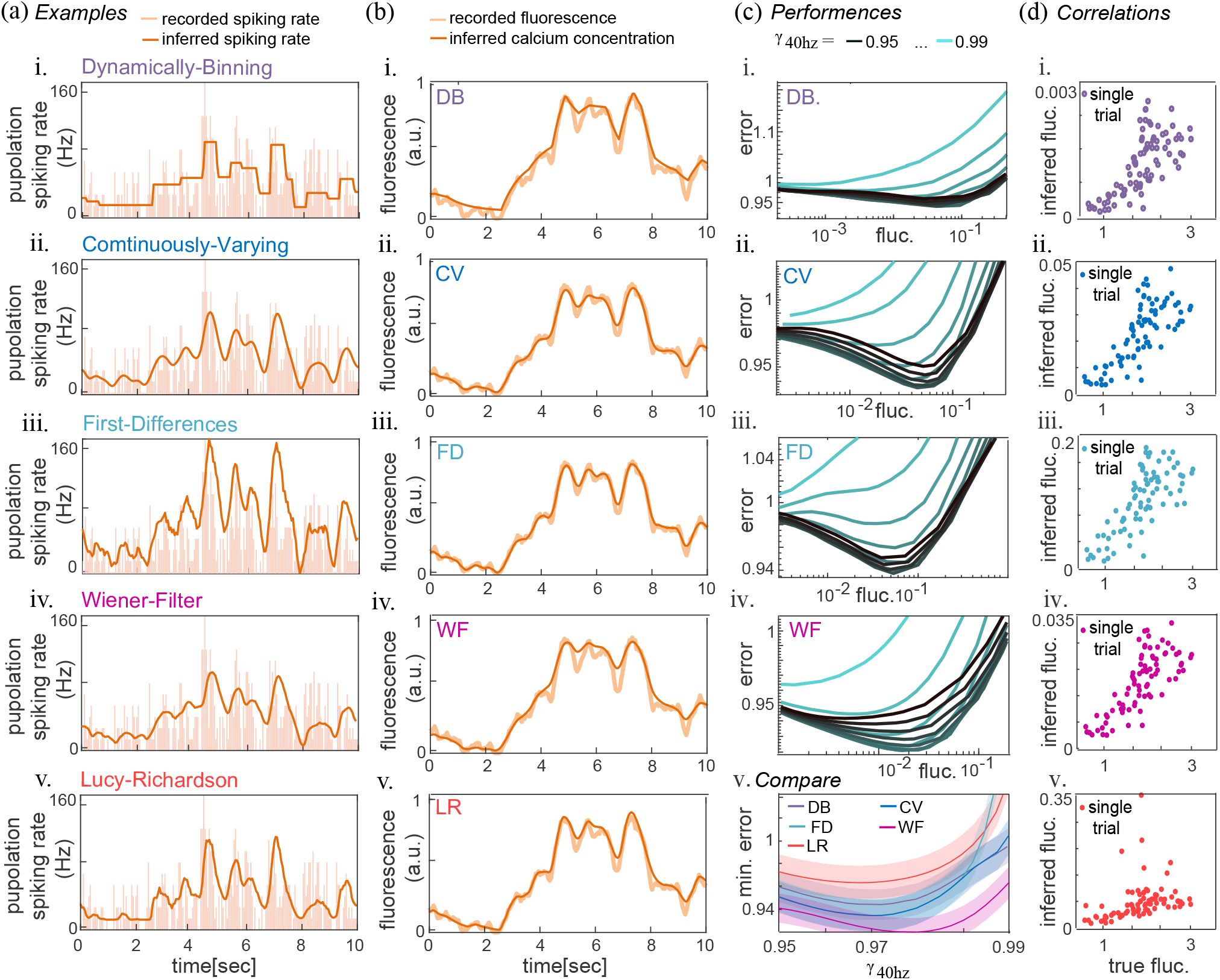
Deconvolution of Fluorescence with Simultaneous Spike Count Recordings. Caption of Figure 3. Dataset no. 1 was segmented into 80 traces for this figure (see Sup. information S3 for details). (a) An example of spike counts recording (ivory bars) displaying a 10-second window from one 20-second segment (b) its simultaneous fluorescence recording (ivory line). (a) The spiking rate (orange line) inferred from the fluorescence trace and (b) the inferred calcium trace (orange line), calculated according to the model (3), from the inferred spiking rate by i. DB, ii. CV, iii. FD, iv. WF, and v. LR. (c)i-iv. The mean error, Equation (18), between the spike counts and the inferred spiking rate for each of the deconvolution methods. Each line corresponds to a deconvolution with a different calcium decay *γ*. The variations in the inferred spiking rate fluctuations, Equation (19), are due to using different parameter values (penalty *λ* in i. DB and ii. CV, different smoothing in iii. FD, or inverse SNR *k* in iv. WF). (c)v. Comparison of the minimum error achieved by the deconvolution methods, as a function of the different decay values *γ*. The minimal mean error for each method is represented by its corresponding colored line, and the standard deviation is indicated with the same colored shade. CV (blue line) yields the second-lowest error, the same as FD (light blue line behind CV). WF (magenta line) yields the lowest error while LR (red line) yields the highest error. (d) The fluctuation level of the inferred spiking rate by i. DB, ii. CV, iii. FD, iv. WF and v. LR, as a function of the spike count fluctuations of each data segment (a single dot).

Finally, the findings in this section agree with the findings using simulations (Sections 3.11 and 3.12) and calcium-only recordings (Section 3.14). By having a partial knowledge about the underlying spiking rates, they establish a needed link between fully controlled, known underlying spiking rates (simulated) data and no information about the underlying spiking rate, as is typical of mesoscale calcium recording experiments.

To evaluate the performance of our deconvolution methods on the datasets from Clancy et al. (2019), we began with a preprocessing analysis of all datasets (details are in Sup. information S3). Following this, we applied our deconvolution methods, Dynamically-Binning, Continuously-Varying, First-Differences, and Wiener-Filter to the preprocessed fluorescence data. We compared the results of these methods with those of the Lucy-Richardson algorithm.

We tested the deconvolution methods using varying calcium decay rates and a range of parameter values (penalty *λ* in Dynbin and Convar, different smoothing in Firdif, or inverse SNR *k* in Wiener). We calculated the error, Equation (18), between the spike counts (recorded alongside the fluorescence, representing the underlying spiking rate) and the inferred spiking rate (generated by each deconvolution method). The different calcium decay rates *γ* produced distinct error curves, while the variation of parameter values (*λ*, smoothing, or *k*) resulted in different inferred fluctuations (Figure 3(c)i-iv).

As evident in the examples (Figure 3(a)), all methods captured the spike counts (ivory bars) by the inferred spiking rates (orange lines) with varying degrees of accuracy. We examined the minimal error achieved at each decay rate (the lowest point in each curve in Figure 3(c)i-iv) across all of our methods. We compared it with the error produced by the Lucy algorithm with the same decay rate (Figure 3(c)v).

For dataset no. 1 in Figure 3, Wiener achieved the lowest error, Convar the second lowest, closely followed by Firdif (most of their curves overlap, as seen in Figure 3(c)v). Dynbin ranked fourth, while deconvolution using Lucy resulted in the highest error.

At the minimal error point, we also calculated the correlation between spike count fluctuations and inferred spike rate fluctuations (19). As explained in section 3.11, a deconvolution method should maintain fluctuation levels after inference. Figure 3(d)i-v displays for each method its inferred spiking rate fluctuations as a function of the spike count fluctuations, for each segment of the data. Wiener maintained fluctuations best, achieving a Pearson correlation of 0.76, followed very closely by Convar with a correlation of 0.75, Firdif with a correlation of 0.72, and Dynbin with a correlation of 0.71. Lucy demonstrated the lowest correlation of 0.32.

We repeated the analysis performed on dataset no. 1 across all Clancy et al. (2019) datasets. The results were qualitatively similar. To better quantify them, we calculated the relative error between the inference errors of our methods and the errors achieved by Lucy (Figure 4(a)). We choose this approach because the magnitude of the spiking activity varies significantly between the different datasets (see the total number of spikes in Sup. information S3 Table 2). And we found that the errors (of all methods) depend linearly on the total number of spikes counted. However, the relative error between one method and another did not exhibit such dependency, nor was it correlated with any dataset property we examined.

**Figure 4:**
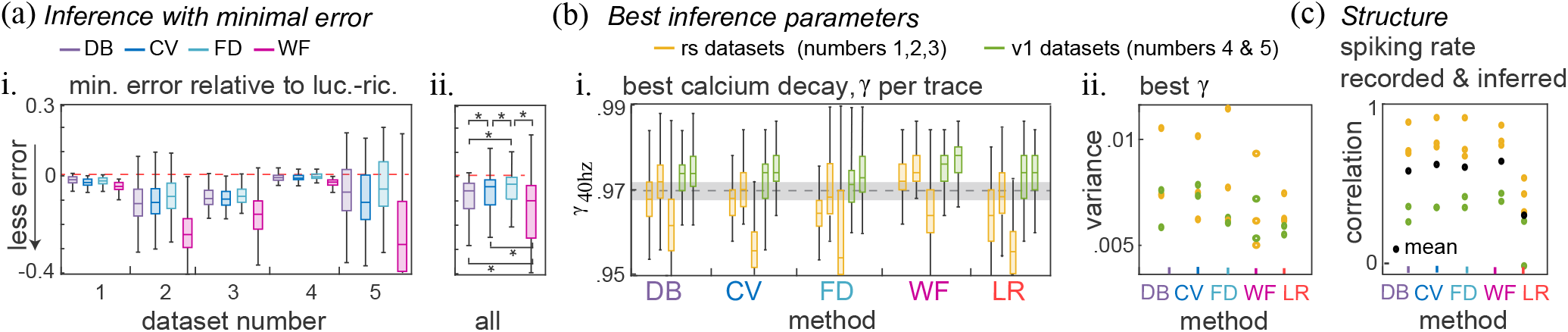
Summary of Deconvolution Results for Simultaneous Fluorescence and Spike Count Recordings. (a)i. The medians (lines inside boxes), quartiles (boxes), and outliers (whiskers) of minimal error achieved by each of our deconvolution methods relative to the minimal error achieved by LR in each of the datasets. The dashed red line marks equal errors (zero difference). Below the red line, the medians of all our methods, DB (purple), CV (blue), FD (light blue), and WF (magenta), performed better than LR across all datasets. ii. For aggregated results across all datasets, the differences between performances were significant. All our methods performed better than LR, with WF performing best, DB second, CV third, and FD last. (b)i. The medians (lines inside boxes), quartiles (boxes), and outliers (whiskers) of the calcium decay with which the minimal error was achieved, for each of the methods (along the *x*-axis) and each dataset (recordings from retrosplenial cortex in yellow and v1 cortex in green). The known calcium decay for single neurons is given by the dashed black line. Note that the dataset identity had more influence on the best *γ* than the deconvolution method. (b)ii. The variance (across segments in each dataset) of the calcium decay with which the minimal error was achieved. Each dataset is marked by a colored dot. (c) Pearson correlations between the spike count fluctuations and the fluctuations of the spiking rate inferred by each of the deconvolution methods. The correlation found for each dataset (by each method) is marked by a colored dot.

**Table 2:** Characteristics of the five parallel fluorescence and spike-count recordings by Clancy et al. 2019.

| Dataset<br>number | Mouse<br>number | Cortical<br>area | Total<br>time (min.) | Percentage<br>analyzed | Total<br># spikes | Fluctuation<br>correlations |
| --- | --- | --- | --- | --- | --- | --- |
| 1 | t41 | rs | 55 | 100 | 195506 | 0.76 |
| 2 | t61 | rs | 60 | 91 | 186765 | 0.90 |
| 3 | t65 | rs | 50 | 87 | 487261 | 0.78 |
| 4 | t42 | V1 | 40 | 96 | 105989 | 0.49 |
| 5 | t64 | V1 | 55 | 98 | 1336222 | 0.47 |

The results indicate that Wiener achieved the lowest error compared to Lucy across all datasets (Figure 4(a)i). When we analyzed each dataset individually for the second and third best methods, and compared all datasets, the results were inconclusive. To address this, we aggregated the data to assess significance across all datasets. Wiener outperformed all other methods as expected, and Dynbin performed significantly better than Convar. Notably, Lucy had the highest error rate, as all other methods consistently demonstrated lower error values across all datasets.

We tested with which calcium decay rate the deconvolution methods achieve minimal error in each dataset and each segment within a dataset (Figure 4(b)). For single neurons, Clancy reported a calcium decay rate of *γ* = 0.97, with a possible deviation of 10% due to physiological variations between cells and variability in mean spiking rates (Chen et al. 2013). This implies that *γ* = 0.97 *±* 0.03 for a single neuron, at the original 40Hz recording rate. Our test results show that the optimal decay rate value for fluorescence deconvolution recorded by the wide-field method was more dependent on the dataset analyzed (and the brain area recorded) than the method itself used for deconvolution (Figure 4(b)i). The median best decay rates, plotted as lines in boxes, are more similar across methods for the same dataset than across datasets inferred with the same method. Crucially, the mean of the best decay rates for inference, considering all datasets, falls within the known decay rate for single cells. Dynbin best decay rate yields 0.97 *±* 0.004, Convar yields 0.969 *±* 0.006, Firdif 0.968 *±* 0.005, Lucy 0.967 *±* 0.007, and Wiener 0.973 *±* 0.005. This suggests that the calcium decay rate for aggregated activity from many single cells can be effectively described using the known decay rate for individual cells, without strong contamination of these values from factors such as glial cells or blood activity. This is important because, in general, only a fluorescence signal is available in mesoscale recordings. Hence, determining the decay rate by comparing the fluorescence signal with parallel spike-count recordings is typically not possible. However, we still need a calcium decay rate for the deconvolution methods. This shows that we can use the known values (for various indicators) from single cells.

We repeated the calculation of the correlation between spike count fluctuations and inferred spike rate fluctuations (Figure 3(d)) across all datasets (Figure 4(c)). The methods Wiener, Convar, and Firdif showed the highest average correlations, with Pearson correlation coefficients of 0.63 *±* 0.22, 0.62 *±* 0.26, and 0.62 *±* 0.24, respectively. This was followed by Dynbin, which had a correlation of 0.57 *±* 0.29. Lucy demonstrated the lowest correlation, with a value of 0.3 *±* 0.22.

### 3.14 Cortex Dorsal Surface Recordings

Here, we evaluate the performance of the deconvolution methods on two “typical” wide-field imaging datasets that include solely fluorescence recordings from the entire dorsal surface of the cortex. The data was recorded by Simon Musall in Anne Churchland’s lab; see Musall et al. (2018) for details. Each of the datasets includes fluorescence recordings from 10 different locations in the cortex (shown in Figure 5(a)i) during 395 trials, each lasting one minute. The recordings were done *in vivo* using GCaMP6s (dataset 1) or GCaMP6f (dataset 2) mice. Dimensionality reduction, subtraction of blood-flow (hemodynamics) related signal, and Δ*F/F* transformation were performed on the traces (Musall et al. 2018) prior to our analysis.

**Figure 5:**
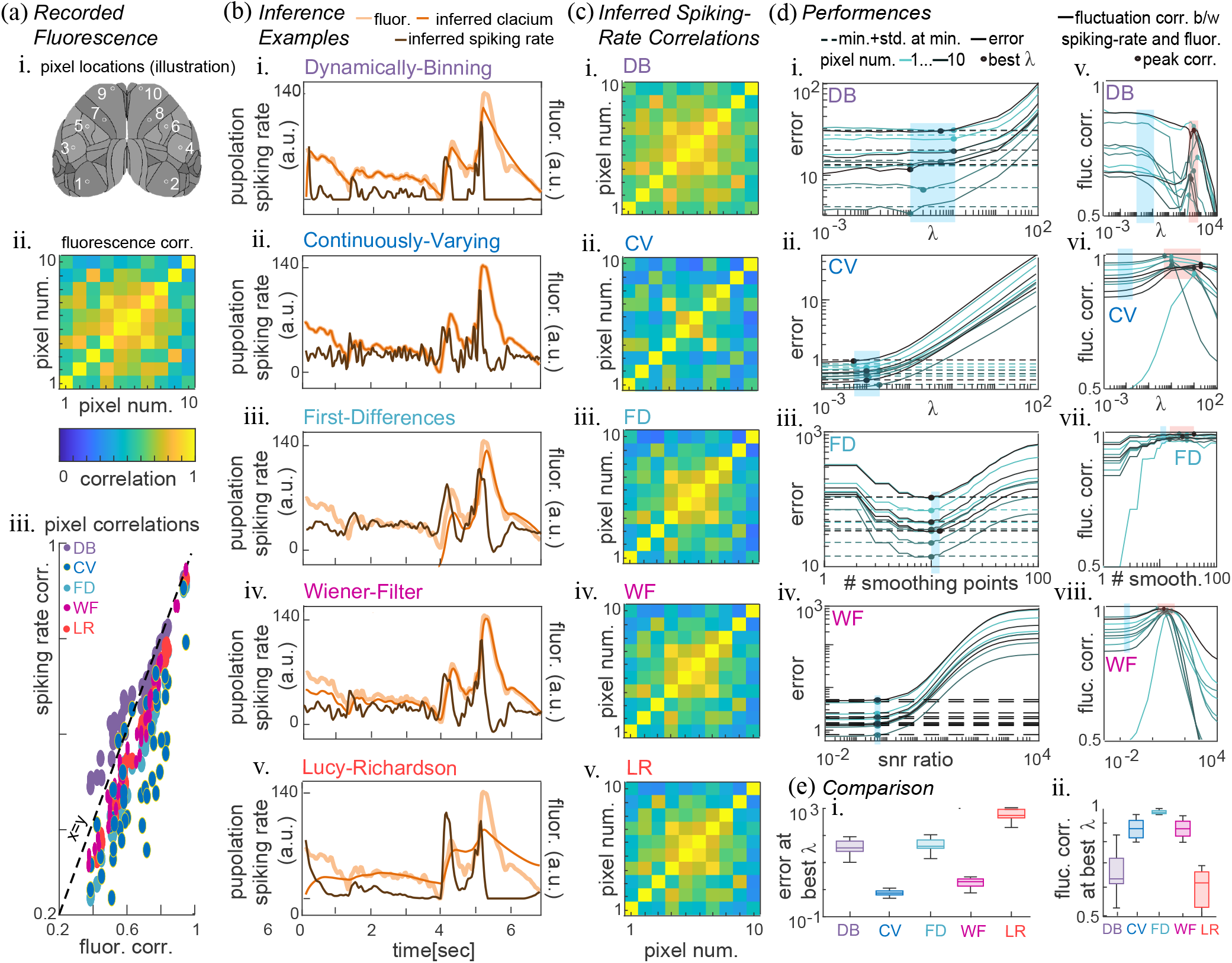
Deconvolution of Cortex-Wide Fluorescence Recordings. Caption of Figure 5 (a)i. Schematic of a whole dorsal surface wide-field view. Numbered white circles indicate the recorded fluorescence locations. ii. Pearson correlations between recorded fluorescence from locations in i. (a)iii. Pearson correlations between all pairs of spiking rate inferred in the ten locations by deconvolution with DB (purple circles), CV (blue circles), FD (light blue circles), WF (magenta circles), or LR (red circles) as a function of their fluorescence Pearson correlations. (b) An example of fluorescence recording (ivory line) and the spiking rate inferred from it (brown line) by i. DB, ii. CV, iii. FD, iv. WF, and v. LR. The calcium concentration, calculated from the inferred spiking rate using the model (3) for each deconvolution method, is plotted in orange. (c) Pearson correlations between all pairs of locations’ inferred spiking rate deconvolved by i. DB, ii. CV, iii. FD, iv. WF, or v. LR. (d)i-iv. The mean error, Equation (20), between fluorescence at odd or even timepoints and inferred calcium from fluorescence at even or odd timepoints, respectively (see algorithm 3 for details). Deconvolution was performed by i. DB, ii. CV, iii. FD, iv. WF. Each line corresponds to one of the ten fluorescence origin locations (each location in a different color, locations are shown in (a)i). Minimal error plus standard deviation at minimal error are marked by a dashed line (same color for each location as the error line). The chosen parameter (penalty *λ* for DB and CV, number of smoothing points for FD, or inverse SNR *k* for WF) by algorithm 3 is marked by a bold dot (same color as its location’s error line). The span of chosen parameters for inference and the corresponding resulting error span is colored in light blue. (d)v-viii. The correlation between fluorescence fluctuations and the fluctuations of the inferred spiking rate by each of the deconvolution methods v. DB, vi. CV, vii. FD, viii. WF. Each line corresponds to one of the 10 fluorescence origin locations (each location in a different color, locations are shown in (a)i). A bold dot marks the local maximum correlation. The span of parameters that maximize the correlations is colored in light pink. For comparison, the span of chosen parameters for inference (d)i-iv. and the corresponding correlation span is colored in light blue. (e)i. The medians (lines inside boxes), quartiles (boxes), and outliers (whiskers) of the ten locations’ inference error by each of our deconvolution methods (y-axis values of bold dots in (d)i-iv) and LR. ii. The medians (lines inside boxes), quartiles (boxes), and outliers (whiskers) of the ten locations’ fluctuation correlations by each of our deconvolution methods (*y* − *axis* values of correlations in (d)v-viii. at the penalty, smoothing, or inverse SNR chosen for inference) and LR.

In Section 3.13, we demonstrated that the calcium decay rate of a single neuron serves as a reliable estimate of the calcium decay rate of the aggregated neural activity. We rely on this finding for analyzing fluorescence-only recordings. Specifically, in this section, we use the known calcium decay rates from single neurons: *γ* = 0.983 for GCaMP6s mice and *γ* = 0.960 for GCaMPsf mice at a recording frequency of 30Hz (Chen et al. 2013).

Since only fluorescence data are available in these datasets (as is common in most mesoscale recordings), we utilize the fluorescence itself as both the training and test sets. We accomplish this by splitting each fluorescence trace 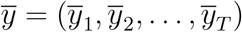 into traces of “odd” indexed timesteps 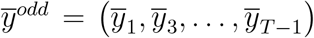 and “even” indexed timesteps 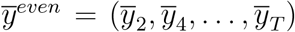 traces. We then infer the “odd” (or “even”) spiking rate from the corresponding “odd” (or “even”) fluorescence using one of the deconvolution methods (with an adjusted calcium decay rate *γ*^2^ that accounts for the trace partition). We compute the mean of the errors of the differences between the “even” or “odd” fluorescence and the “odd” or “even” inferred calcium, respectively. Where the “odd” (or “even”) inferred calcium is calculated from the “odd” (or “even”) inferred spiking rate using our model (3). The resulting error is given by

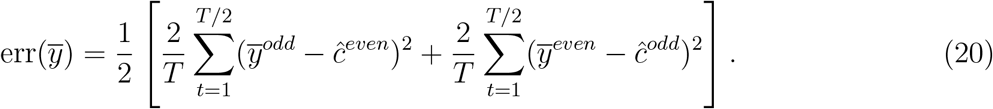

Changing the penalty value in Dynbin and Convar, adjusting the smoothing in Firdif, or modifying the inverse SNR *k* in Wiener affects the error, Equation (20). We chose to infer with the maximal penalty (smoothing, or inverse SNR) value that keeps the error below its minimal value plus its standard deviation (across trials) at the penalty value with minimal error. A complete description of this procedure can be found in Algorithm 3 in Sup. information S4. This method was introduced by Jewell et al. (2019) (note that *ycMSE* denotes the error in (20)).

While the magnitude of the error, Equation (20), varies between different pixels (see Figure 5(d)i-iv), the shape of the error curve remains consistent. The optimal parameter (penalty, smoothing, or SNR) for deconvolution remains similar across cortical regions due to this consistency. This further indicates that these error curves are shaped by the characteristics of neural and calcium dynamics, which are naturally shared between regions. In contrast, the magnitude of luminescence can vary across cortical areas depending on their physical distance from the camera. However, this primarily affects the magnitude of the error and does not impact the choice of deconvolution parameters.

Regarding the performance of the deconvolution methods, both per pixel and on average, Convar produced the least error, Wiener the second lowest, and Lucy the largest error, while Dynbin and Firdif showed intermediate values, see Figure 5(e)i. In previous sections, we demonstrated that effective deconvolution preserves the data structure, i.e., yields high correlations between fluctuations of the fluorescence traces and those of the inferred spiking rates. With the optimal penalty smoothing or SNR, Firdif maintained the highest correlations, Convar and Wiener showed the second highest, while Dynbin performed slightly better than Lucy, which had the lowest correlations; see Figure 5(e)ii.

Another measure of deconvolution performance is its ability to remove “ghost” correlations between pixel pairs caused by the calcium dynamics. The slow decay of the calcium levels can lead to “ghost” correlations—these are excessive correlations that do not exist to that extent in the underlying neural dynamics. They occur because calcium levels remain elevated even after the neural activity that triggered the increase has ceased. During these times, the elevated calcium activity from one pixel can correlate with a different calcium-driven neural activity in another pixel, even if those neural-driven activities do not originally correlate with one another.

As a result, when analyzing recorded fluorescence data, most pixel-to-pixel correlations tend to decrease with the inferred spiking rates obtained through a deconvolution method, see Figure 5(a)iii. Notably, Convar not only reduces these ghost correlations on average but also differentiates between various pixel pairs, decreasing correlations by different percentages depending on the specific pair (see Figure 5(a)iii blue circles; this is also visual through the differences between the functional correlation matrices using the fluorescence in Figure 5(a)ii and using the inferred spiking rates by Convar in Figure 5(c)ii). It is likely that it effectively distinguishes between correlations that appear high due to ghost effects and those that are genuinely high due to underlying neural activity. Wiener, Firdif and Lucy also successfully reduce ghost correlations by different percentages, to some extent. In contrast, Dynbin not only fails to reduce these correlations but also produces spike rates with high correlations in pixel pairs that exhibit lower correlations in their fluorescence data.

The analysis presented above is supported by examining the pixel-to-pixel correlation with GCaMP6f data. If ghost correlations arise due to the slow decay of calcium, GCaMP6f mice should exhibit fewer ghost correlations compared to GCaMP6s mice, as the calcium signal that contributes to these unrealistic correlations decays more quickly in GCaMP6f mice (assuming all other conditions, including the experimental setup, are kept constant). A simple visual examination confirms it. The functional correlations matrix calculated from the fluorescence data of GCaMP6s (Figure 5(a)ii) contains higher correlation values (indicated by yellow and orange squares) than the functional correlations matrix generated with GCaMP6f fluorescence data (Figure 6(a)i). Accordingly, the various deconvolution methods were successful in reducing excessive correlations in GCaMP6f (Figure 6(a)ii), although the changes in values were less pronounced than in GCaMP6s.

**Figure 6:**
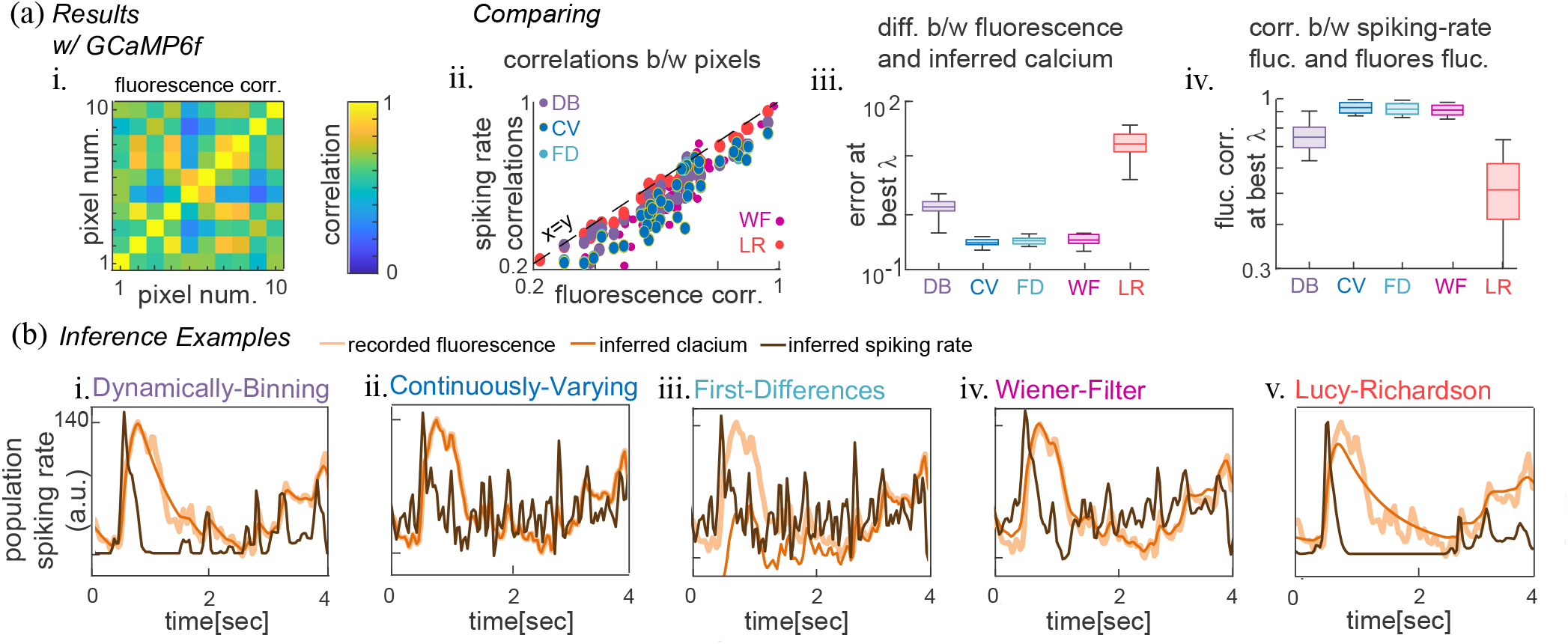
Summary of Cortex-Wide Deconvolution for Recordings with GCaMP6f. (a)i. Pearson correlations between recorded fluorescence traces in different locations (locations are illustrated in Figure 5(a)i). ii. Pearson correlations between all pairs of spiking rate inferred in the ten locations by deconvolution with DB (purple circles), DV (blue circles), FD (light blue circles), WF (magenta circles), or LR (red circles) as a function of their fluorescence Pearson correlations (shown in (a)i). iii. The medians (lines inside boxes), quartiles (boxes), and outliers (whiskers) of the ten locations’ inference error between odd or even fluorescence timepoints and inferred calcium from even or odd fluorescence timepoints, respectively, for inference with DB (purple), CV (blue), FD (light blue), WF (magenta), and LR (red). iv. The medians (lines inside boxes), quartiles (boxes), and outliers (whiskers) of the ten locations’ fluorescence and inferred spiking rate fluctuation correlations by each of the deconvolution methods: DB (purple), CV (blue), FD (light blue), WF (magenta), and LR (red), at the chosen penalty, smoothing or SNR, for inference. (b) An example of fluorescence recording (ivory line), the spiking rate inferred from fluorescence (brown line) by i. DB, ii. CV, iii. FD, iv. WF, and v. LR. The calcium concentration, calculated from the inferred spiking rate using the model (3), is plotted in orange.

Overall, the various deconvolution methods demonstrate similar performance on GCaMP6f fluorescence data (Figure 6) as they do on GCaMP6s fluorescence data (Figure 5). One notable difference is the ability of Firdif to produce low error, Equation (20), with the GCaMP6f data and performs as well as Convar (Figure 6(a)iii). The fluctuation correlations between the fluorescence traces and the spiking rates inferred by Firdif and Convar are similar with GCaMP6f data, achieving both best results (Figure 6(a)iv). This manifests in the inferred spike rate fluctuations, as visually evident by the rapidly changing inferred spiking rate examples provided for both Convar and Firdif for GCaMP6f data (Figure 6(b)ii and iii, brown lines). The lower error and higher correlations achieved by Convar and Firdif (and Wiener to some extent) suggest that their high inferred spiking rate fluctuations (compared to Firdif and Lucy) accurately reflect actual underlying neural activity rather than noise. In contrast, Dynbin and Lucy tend to be oversmooth during inference, overlooking the fluctuations present in the neural activity. The performance of Convar on GCaMP6s reinforces this observation, as its example also reveals a highly fluctuating inferred spiking rate (Figure 5(b)ii) compared to other methods while achieving the best results in terms of minimal error and pixel-to-pixel correlation differentiation.

### 3.15 Hemodynamics Impact

The wide field of view and relatively low resolution of mesoscale imaging can cause calcium fluorescence measurements of neural activity to be contaminated by light emitted from nearby blood vessels (Gilad 2024). Because blood fluorescence intensity varies with blood volume, these light signals can interfere with the recorded neural activity. To remove them from the recording, an additional light source insensitive to calcium fluorescence emission (typically green or ultraviolet light) is recorded in interleaved sequences with the regular excitation light (blue light at 473 nm for GCaMP6) (Ma et al. 2016). The fluorescence associated with neural activity is then determined by subtracting the signal from the non-calcium-emitting fluorescent light (green or ultraviolet) from the total recorded light (blue). An example is shown in Figure 7(a)i-iii..

**Figure 7:**
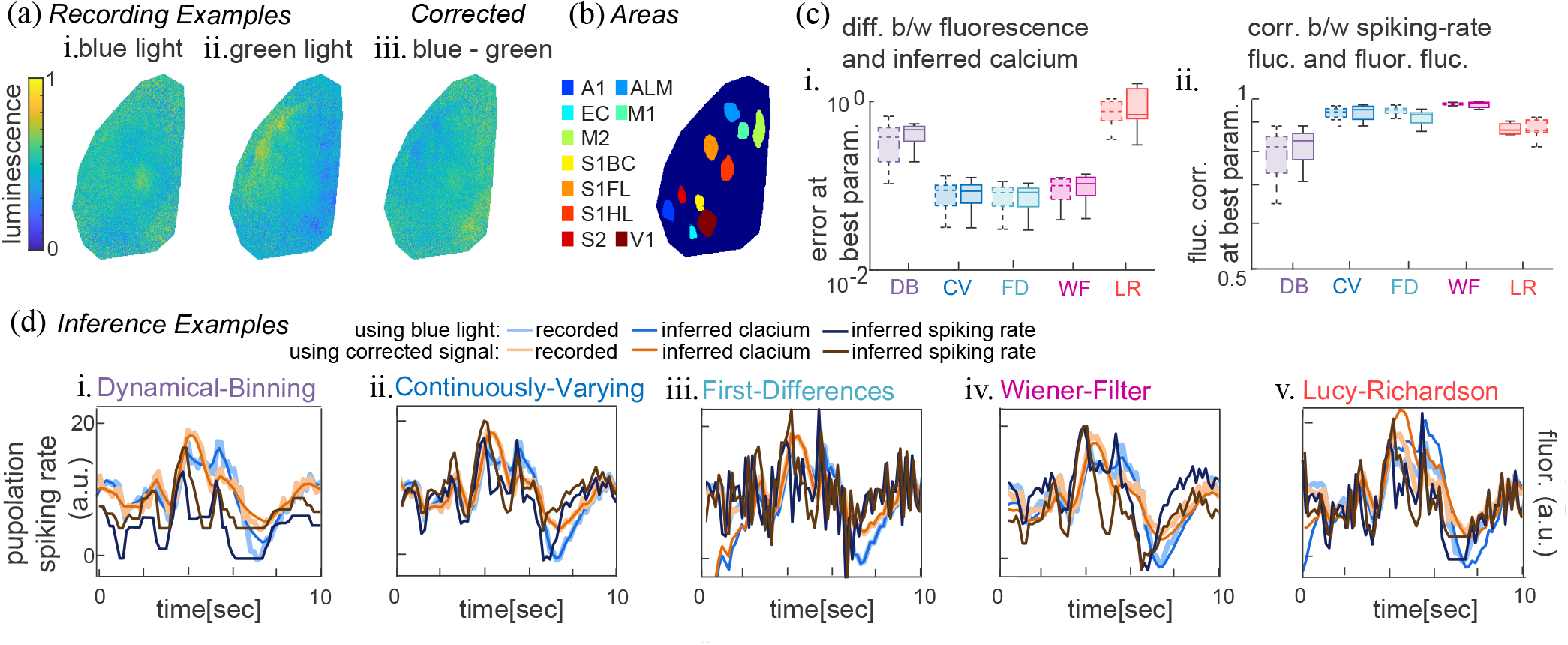
Hemodynamics and Deconvolution. (a) An example of wide-field fluorescence imaging at a specific timepoint. Intensities of light recorded across the dorsal surface of the left hemisphere. i. Captured blue light (total signal) ii. Captured green light (hemodynamics signal) iii. The intensity of the captured blue light minus the intensity of the green light (neural activity signal). (b) locations of the brain areas analyzed, as identified by Gilad et al. 2018. Each area’s average blue light intensity (neural activity and hemodynamics) and blue light minus green light intensity (neural activity with blood-flow luminescence subtracted) were deconvolved. Areas include: auditory areas (A1), entorhinal cortex (EC), anterior lateral motor cortex whisker-related (ALM), primary motor cortex (M1 and M2), barrel cortex in primary somatosensory cortex (S1BC), rostro-lateral cortex corresponding to the whisker evoked activation (S1FL and S1HL), secondary somatosensory cortex (S2), and primary visual cortex (V1). (c)i. The medians (lines inside boxes), quartiles (boxes), and outliers (whiskers) of the ten areas’ inference error, Equation (20), achieved by each of the deconvolution methods inferring the blue light recorded signal (dashed boundary lines) and the corrected signal after blood-flow luminescence subtraction (solid boundary lines). ii. The medians (lines inside boxes), quartiles (boxes), and outliers (whiskers) of the ten areas’ fluctuation correlations at the best parameter value chosen for inference, achieved by each of the deconvolution methods inferring the blue light recorded signal (dashed boundary lines) and the corrected signal after blood-flow luminescence subtraction (solid boundary lines). (d) For each of the deconvolution methods i. DB, ii. CV iii. FD, iv. WF and v. LR, the following is shown: an example of corrected fluorescence recording (ivory line), the spiking rate inferred from it by the deconvolution method (brown line), and the calcium concentration calculated according to the model (3) from the inferred spiking rate (orange line). The blue light (total activity) signal recorded in parallel (light blue line), the spiking rate inferred from it by the deconvolution method (dark blue line), and the calcium concentration calculated according to the model (3) from the inferred spiking rate (blue line).

This study examines the impact of blood-flow related signal subtraction on the temporal deconvolution of recorded fluorescence signals. Such substruction alters the fluorescence signal, which in turn affects the inferred spiking rate, as illustrated in Figure 7(d). Excluding the blood fluorescence signal from the total recorded luminescence results in corresponding changes to the inferred spiking rate. For the deconvolution process to be robust, its performance and parameter selection should remain unaffected when blood-flow-related signals are removed from the fluorescence data. The expectation is that the deconvolution process, including the evaluation of optimal penalty (smoothing or SNR), the measurement of error, and the assessment of fluctuation correlations, remains consistent regardless of the presence or absence of blood-flow signals. This consistency is essential to ensure that blood flow obstruction does not introduce unintended effects on the deconvolution process beyond the desired signal shifts.

We tested a wide-field imaging dataset recorded by Ariel Gilad in the lab of Fritjof Helmchen (Gilad et al. 2018) before and after blood-flow luminescence signal subtraction. Our results indicate that removing blood-flow-related luminescence does not affect the deconvolution process for any of the tested methods. Measurements of error (Figure 7(c)i, dashed versus solid boundary lines) and correlations of fluctuations (Figure 7(c)ii, dashed versus solid boundary lines) show no significant differences regardless of whether blood-flow luminescence was removed. Furthermore, the optimal parameter values (penalty, number of smoothing points, or inverse SNR) remained unchanged.

Our analysis of the optimal calcium decay rate in Section 3.13 supports this conclusion. It showed that the calcium decay rate of a single neuron closely aligned with the calcium decay rate observed in fluorescence recordings from mesoscale imaging. Because fluorescence emission from blood vessels does not decay in the same manner as calcium in response to neural activity, our findings suggest that contamination from nearby light sources, including blood vessel fluorescence, has a limited impact on wide-field recordings of neural activity and does not significantly obscure or alter the recorded neural dynamics.

## 4 Discussion

The problem of inferring spiking rates from fluorescence recordings that capture aggregate neural activity presents several challenges. One such challenge is the absence of experiments that record in parallel the underlying neural activity and the fluorescence, providing ground truth examples of the spiking rate that generates the fluorescence. It is missing simply because a technology that allows such recordings does not exist. The absence of such parallel recordings impedes the training of convolutional deep networks or similar blind neural network training approaches for spiking rate inference from mesoscale fluorescence imaging.

Existing signal processing techniques, such as the Lucy-Richardson algorithm and the Wiener-Filter method, which we have adapted for temporal spiking rate inference and evaluated herein, rely on assumptions that do not necessarily match the structure of mesoscale imaging data. In particular, the Lucy-Richardson algorithm assumes Poisson noise, which is appropriate for blurred images but less suitable for mesoscale fluorescence recordings. In these recordings, noise can originate from multiple sources, including recording equipment, light reflections, and animal movement, and is unlikely to follow Poisson statistics. This mismatch possibly contributes to the reduced performance of the Lucy-Richardson algorithm, including its difficulty in distinguishing signal from noise and hence in recovering the true range of underlying spiking rates. By contrast, the Wiener Filter method performs competitively in some of the real recording settings. The Wiener Filter assumes a fixed relationship between signal and noise, which is possibly a reasonable approximation in fluorescence recordings, where both signal and noise can scale with luminescence magnitude.

The two novel algorithms we introduce here, the Dynamically-Binning algorithm and the Continuously-Varying algorithm, are designed to impose minimal structural constraints required to recover spiking rates from mesoscale fluorescence data, including non-negativity of the inferred spiking rates and temporal consistency with calcium dynamics. Thus, their improved performance likely reflects the fact that their assumptions better match mesoscale calcium imaging than those underlying standard signal-processing deconvolution methods. We prove that the optimization problem solved by the Dynamically-Binning algorithm and the Continuously-Varying algorithm is convex, meaning they reach, within their respective constraints, optimal solutions with minimal error.

We conducted a rigorous evaluation of the various inference approaches we developed, employing multiple datasets and a range of performance metrics. The datasets included simulations, parallel recordings of fluorescence and localized spike counts, as well as fluorescence-only data. To facilitate a comprehensive understanding of the comparative performance of the different methods, we present a summary in Table 1. For each dataset and metric, the table assigns each method to one of four qualitative performance categories: Excellent, Good, Fair, or Poor. Because several methods can perform similarly well on a given metric, more than one method could be assigned the same category. We also included their relative performance in terms of computational demands(run time)^3^.

Prior to this study, Lucy-Richardson was the only published deconvolution method for the analysis of mesoscale calcium recordings (Wekselblatt et al. 2016). In this study, we introduce four alternative approaches that outperformed Lucy-Richardson across all datasets and most measurements. While we evaluated additional methods beyond those presented here, we surveyed in this work only those we found fit (i.e., performed better than Lucy-Richardson in most cases).

Our findings indicate that Continuously-Varying is the most suitable approach across diverse datasets, measurements, and conditions. Although it did not consistently outperform other methods in every scenario, it always performed well. In most scenarios, it performed best or second best. Importantly, it never yielded poor results. Notably, the Wiener-Filter method also demonstrated reliable capabilities across varying conditions. As a practical recommendation, we suggest generating error curves for inference by comparing odd and even timepoints within the data (following Algorithm 3). Curves that replicate the structure illustrated in Figure 5(d), exhibiting similar decay patterns across pixels with varying amplitudes, yet providing comparable best penalty values, suggest a good fit of the data to the applied approach. Conversely, when recordings are unstable or affected by reflections, the resultant curves will not yield similar best penalties. The correlations between pixels in this case will also remain high, a discrepancy that also serves as a good measure of the quality of the inference (see Figure 5(a)iii&(c) and Figure 6(a)ii). We recommend considering Wiener-Filter as a secondary option. Since it is based on a different approach, it may be better suited for datasets that Continuously-Varying struggles to capture effectively. In any case, all methods are superior to using the fluorescence as a measure of the spiking rate, as the error between the observed 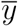 fluorescence and the underlying simulated (or estimated measured) spiking rate *r* is an order of magnitude higher than the largest among the methods’ errors produced by Lucy-Richardson.

Last, when deconvolving, key information becomes more accessible rather than being lost. While the absolute measures of the spiking rates and their baseline activity remain unknown, the same limitations apply to fluorescence data. The use of calcium dynamics (model) to infer the spiking rate incorporates additional information beyond what fluorescence provides. With this information, the correct temporal dynamics of neural activity is gained, and noise is removed according to it. It is imperative to remember that the dynamics of spiking rates are significantly faster, 10 to 100 times, than the dynamics of fluorescence. Consequently, the inferred spiking rates may appear noisy; however, they are, in reality, simply faster yet less noisy thanks to the noise removal that all methods perform. Ultimately, inference results offer a more reliable depiction of the underlying neural dynamics.

## 5 Acknowledgements

We are grateful to all the people who provided data for this study: Kelly Clancy (recorded at Thomas Mrsic-Flogel Lab), Simon Musall (Recorded at Anne Churchlands Lab), and Ariel Gilad (Recorded at Fritjof Helmchen Lab). ESB was supported by NSF DMS Grants 1514743 and CAREER-1056125. We also acknowledge support from the Sackler Foundation, the Swartz Foundation via the Swartz Center for Theoretical Neuroscience at the University of Washington, and the NSF-ERC Center for Sensorimotor Neural Engineering at the University of Washington. We thank The Rockefeller University for its unique support. We thank the Allen Institute founders, Paul G. Allen and Jody Allen, for their vision, encouragement and support.

## Supplemental Information

S1 Theory and Supporting Mathematical Derivations

S2 Simulation Details

S3 Parallel fluorescence and Spike Count Recording Datasets Including Table 2

S4 Calculating the Error and Choosing Penalty *λ* from Solely Fluorescence Recordings Including Algorithm 3

## S1 Theory and Supporting Mathematical Derivations

### S1.1 Proposition 1

#### Proposition 1.

*The optimization problem* (4) *is equivalent to*

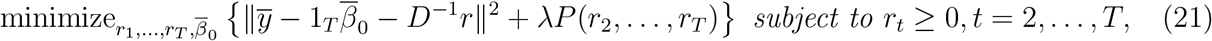

*in the sense that* 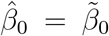 *and* 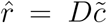, *where* 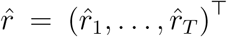 *and* 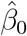 *solve* (21), *and* 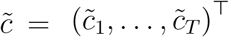 *and* 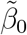 *solve* (4).

The optimization problem (21) with the penalty (5) takes the form of the optimization problem (6).

### S1.2 Proposition 2

We make the following observation to further simplify problem (6):

#### Proposition 2.

*The pair* 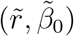 *is a solution to problem* (6) *if and only if* 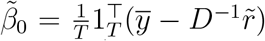 *and* 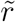 *is a solution to problem* (8) *where* 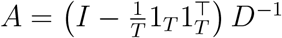 *and* 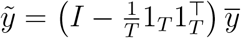, *with I is the T × T identity matrix and* 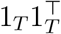 *is a T × T matrix of ones*.

#### Lemma 1.

*Let A* = *PD*^−1^, 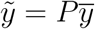 *with* 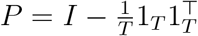 *and* 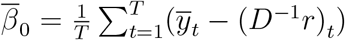. *Then*, 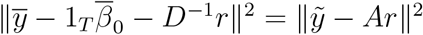.

#### Proof of Lemma 1.

Since *P*^⊤^*P* = *P* and *P* 1_*T*_ = 0, it follows that

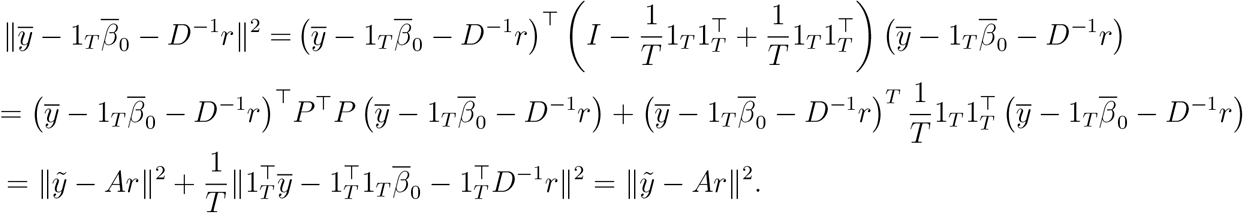

The last equality follows from the fact that 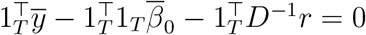.

#### Proof of Proposition 2.

Taking the derivative of (6) with respect to 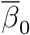 and setting it equal to zero, we find that 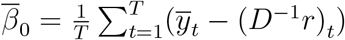. The result follows from Lemma 1.

### S1.3 The Proximal Gradient Descent Algorithm

In summary, the proximal gradient descent solves optimization problems of the form

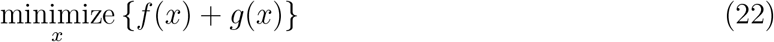

where *f* (·) is a smooth convex function, and *g*(·) is convex but possibly non-differentiable. Then, under mild conditions, an iterative algorithm that initializes *x* at *x*^(0)^, and then at the *t*th iteration applies the update *x*^(*t*)^ ← Prox_*sg*(·)_ *x*^(*t*−1)^ − *s*∇*f* (*x*^(*t*−1)^), will converge to the global optimum. Here, *s* is a stepsize chosen so that *s* ≤ 1*/L*, where *L* is the Lipschitz constant for the function ∇*f* (·). The notation Prox_*sg*(·)_ indicates the *proximal operator* of the function *sg*(·), defined as

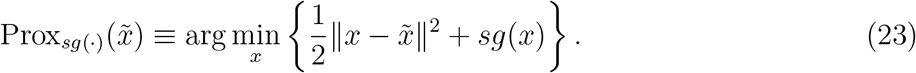

Therefore, proximal gradient descent provides a simple recipe for solving a broad class of convex optimization problems of the form (22), provided that the proximal operator (23) is easily computed, and the function ∇ *f* (·) is Lipschitz continuous. A detailed treatment can be found in Parikh et al. (2014).

### S1.4 Proposition 3

Using the notation of (22), to solve (10), we take 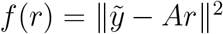 and 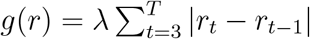. The following result will be useful,

#### Proposition 3.

*The function* 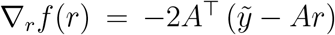 *is Lipschitz continuous with Lipschitz constant L satisfying* 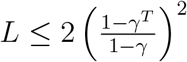.

Note that Equation (3) implies

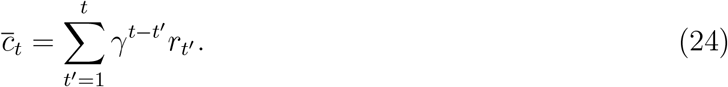

Since 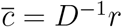, we find from (24) that the entries of *D*^−1^ are given by

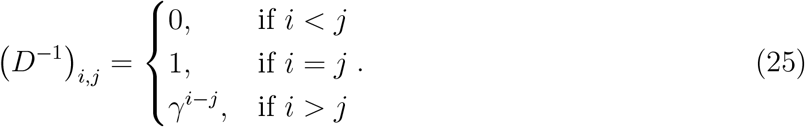

#### Lemma 2.

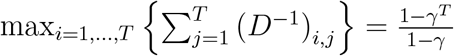.

*Proof*. From (25),

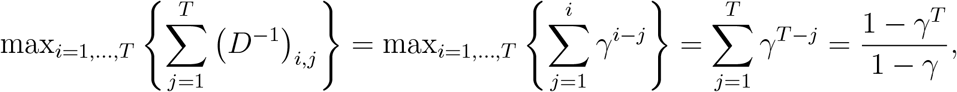

where the last step was calculated using the expression for the sum of a geometric series. Following the same reasoning, one can also show that 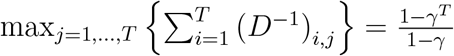.

#### Proof of Proposition 3.

The Lipschitz constant of 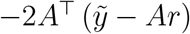 is given by the largest eigenvalue of 2*A*^⊤^*A* (Bubeck et al. 2015). To find it, it is convenient to first explore the largest eigenvalue of (*D*^−1^)^⊤^ *D*^−1^, which is a symmetric matrix with positive real entries. By the Perron Frobenius Theorem and Lemma 2, we can bound its largest eigenvalue by its largest single row sum:

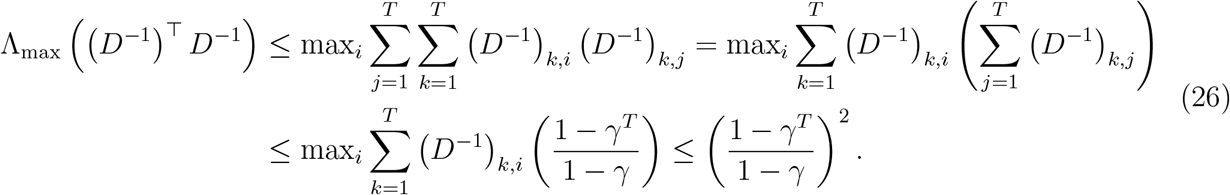

We recall that *A* = *PD*^−1^ and we observe that *P*^⊤^*P* = *P*, where 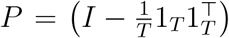, with *I* the identity matrix and 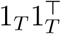 a *T T* matrix of ones. Together with (26) and Rayleigh quotient properties, the following holds for any vector *u*:

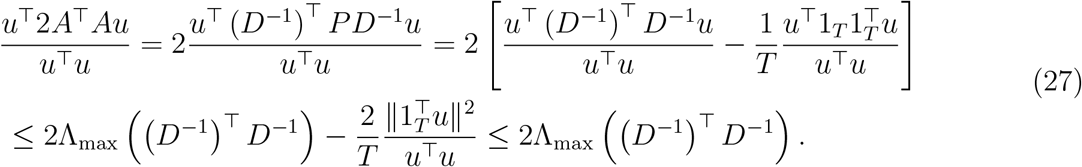

Since the above inequality holds for any vector *u*, including the eigenvector associated with the largest eigenvalue of 2*A*^⊤^*A*, it follows that

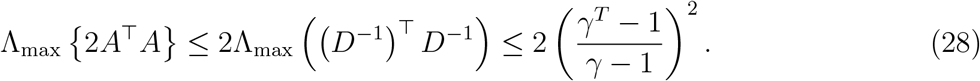

Hence the Lipschitz constant of 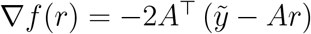 is bounded above by 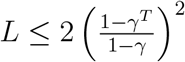.

### S1.5 Proposition 4

#### Proposition 4.

*Let* 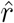 *solve the optimization problem*

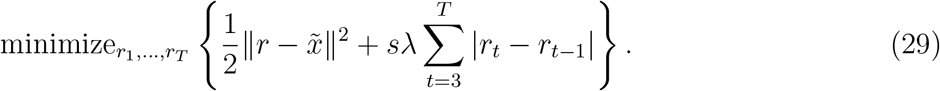

*Then* 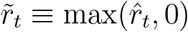 *for t* = 2, …, *T and* 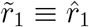 *solves the optimization problem*

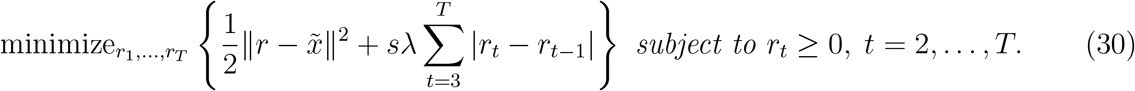

A number of standard solvers, such as the `flsa` solver in `R` (Hoefling 2010), are available to solve (29). We solve (29) by implementing the proposal of Condat (2013). Proposition 4 implies that given a solution to (29), solving (30) is straightforward.

We will make use of the *generalized sign*, the subdifferential of the *ℓ*_1_ norm, defined as

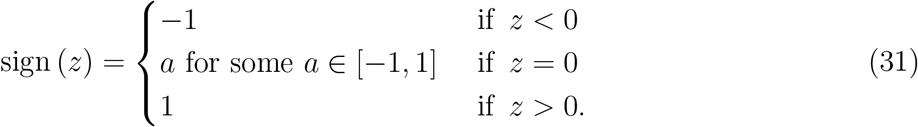

When *z* is a vector, the operation sign(*z*) is applied componentwise.

#### Proof of Proposition 4.

The solution 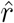 to (29) satisfies the optimality condition

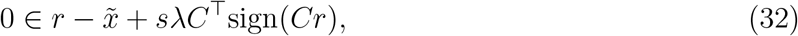

where *C* is a (*T* − 2) *× T* matrix defined as

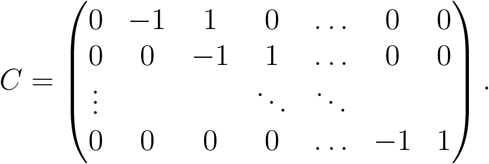

Furthermore, 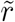 is a solution to (30) if and only if there exists some 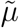 such that the pair 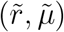 satisfies the Karush-Kuhn-Tucker optimality conditions (Boyd & Vandenberghe 2004), given by

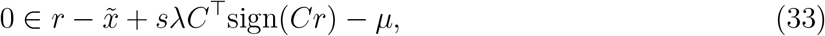

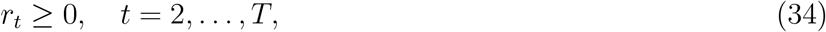

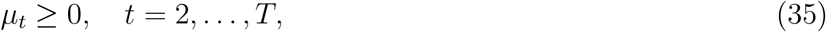

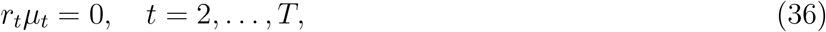

where *µ*_1_ = 0. To complete the proof, we will show that 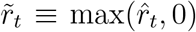 and 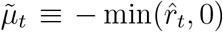 satisfy (33)–(36).

The fact that 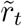 and 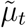 satisfy (34)–(36) follows by inspection. It remains to show that 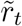 and 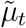 satisfy (33). Notice that 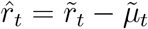. Therefore, because 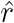 solves (29), it follows directly from (32) that

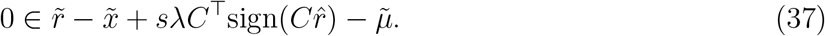

Inspection of the matrix *C* reveals that the elements of 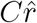 are of the form 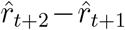. Furthermore, it is straightforward to show that (i) 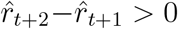 implies 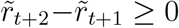; (ii) 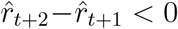 implies 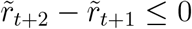; and (iii) 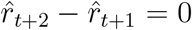 implies 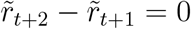. Therefore, 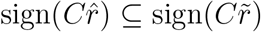. Combining this with (37) directly implies that the pair 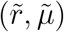 satisfies (33).

Propositions 2, 3, and 4 lead directly to Algorithm 1 for solving (10).

### S1.6 Proposition 5

We can further express (14) as

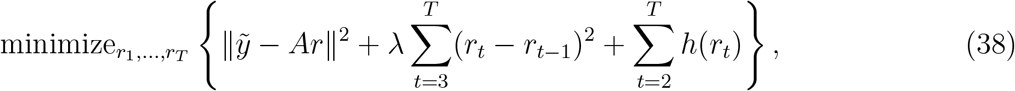

with

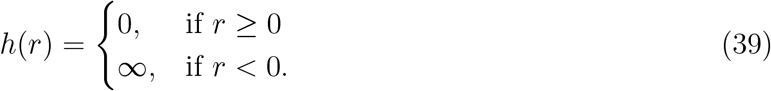

To utilize the proximal gradient decent (see Equations (22) and (23)), we further express the objective function in (38) as *f* (*r*) + *g*(*r*) where

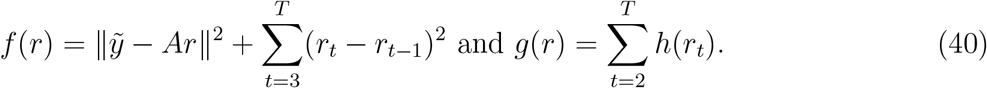

#### Proposition 5.

*The function* 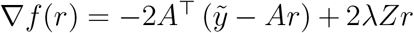 *is Lipschitz continuous with Lipschitz constant L satisfying* 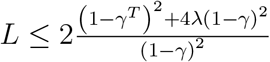, *where Z defined in* (13).

We note that 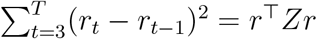.

#### Proof of Proposition 5.

The Lipschitz constant of the function 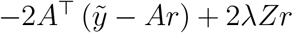 is given by the largest eigenvalue of 2*A*^⊤^*A* + 2*λZ* (Bubeck et al. 2015). Since *A*^⊤^*A* and *Z* are symmetric and real, the largest eigenvalue, Λ_max_, of their sum is bounded by the sum of their largest eigenvalues:

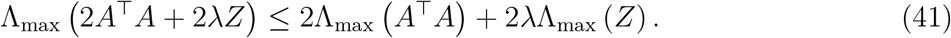

It is not hard to show that Λ_max_ (*Z*) ≤ 4.

Together with (28) and (41), it follows that the Lipschitz constant *L* satisfies 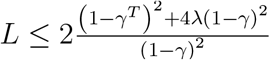.

### S1.7 Proposition 6

#### Proposition 6.

*For any s* ≥ 0, *the solution to the optimization problem*

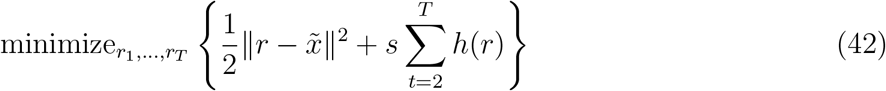

*is given by* 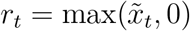 *for t* = 2, …, *T and* 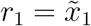.

Proposition 6 is straightforward and the proof is omitted. Propositions 2, 5 and 6 lead to Algorithm 2 for solving (14).

### S1.8 Solutions are Invariant Under a Constant Shift

With either *n* = 1, for ‘Dynamically-Binning’, or *n* = 2, for ‘Continuously-Varying’, (6) is a convex optimization problem. However, it is not strictly convex and so the solution is not unique. The following proposition indicates that the solution to (6) is invariant under a constant shift in the spiking rate.

#### Proposition 7.

*Let the pair* 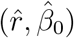 *denote a solution to* (6). *Then the pair* 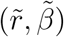 *also solves* (6), *where* 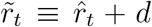 *for t* = 2, …, *T for any* 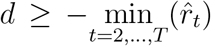 *that satisfies* 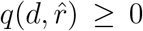, *for* 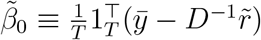 *and a particular choice of* 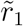.

#### Proof of Proposition 7.

Proposition 2 states that 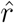 is a solution to (6) if and only if it is also a solution to (8). Hence, 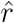 minimizes the objective function 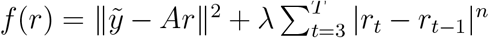. Therefore, any 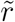 that satisfies 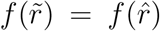 with 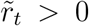 for *t* = 2, …, *T* is a solution to the optimization problem (6) as well.

If we construct 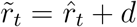 for *t* = 2, …, *T* with 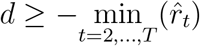, then 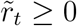 for *t* = 2, …, *T*.

In addition, 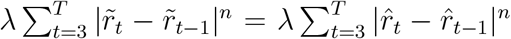. So to show that 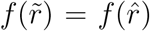, it suffices to show that 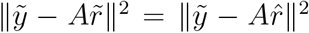. We will do so by choosing an appropriate value for 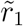. In particular, notice that

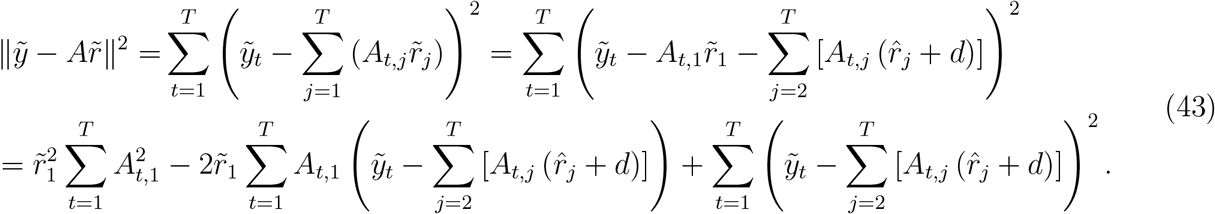

So it suffices to find the value of 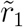 for which (43) equals 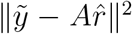. We observe that (43) is quadratic in 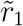. We define

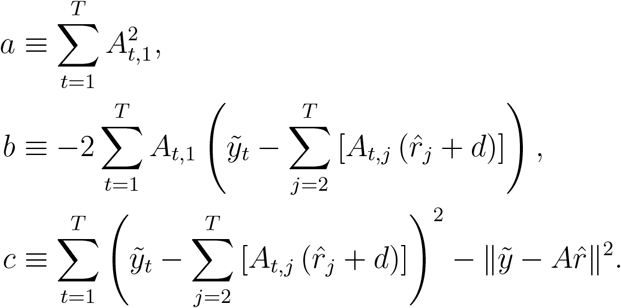

A solution exists provided that *b*^2^ − 4*ac* ≥ 0, in which case 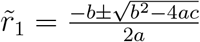.

Defining 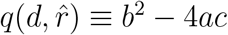, the proof is complete.

## S2 Simulation Details

We generated spiking rate traces *r* by integrating the neural network given by tanh(*x*), where *x* is a vector of size *N* = 2000, *g* = 5.7, *J* is an *N × N* matrix with i.i.d. entries *J*_*ij*_ ~ *N* (0, 1*/N*) and *τ* = 1*sec*. We sampled the activity of *r* = tanh(*x*) for *x* = 1 … 1000 every 50ms for 5sec. We repeated this process with 10 different networks, resulting in 10000 spiking rate traces with 1000 timepoints. To ensure non-negativity of the rate trace, we subtracted out their minimum activity. To change the fluctuations of *r* we run the networks with variable *g*’s.

## S3 Parallel Fluorescence and Spike Count Recording Datasets

We analyze five datasets recorded by Clancy K. in the Mrsic-Flogel lab at UCL. These datasets consist of parallel recordings of spike counts and wide-field fluorescence in vivo, obtained from either the visual cortex (V1) or the retrosplenial cortex (RS) of mice with GCaMP6f (see (Clancy et al. 2019), Supplemental Figure 4). The spike counts were recorded at the exact same physical location from which the fluorescence originated. The table below provides details on the recoded datasets, including the duration of each recording, the percentage of that duration used in our analysis, the total number of spikes counted, and the correlation between the fluctuations in fluorescence and in spike counts for each data set.

Clancy et al. recorded their datasets at 40 Hz. Our first analysis step was to average every two consecutive data points to create a dataset at 20 Hz. This adjustment allows us to work with a bin size of 50ms, which is longer than the calcium rise time of approximately 40ms (Chen et al. 2013). As a result, we can assume that the calcium concentration rises instantly within a bin, while decay occurs over many bins because the decay times are significantly longer. Additionally, averaging the two data points reduces noise, enabling us to more directly compare the deconvolution methods.

Our second analysis step involved estimating *β*_1_ for each dataset. Recall that in the single neuron model (1), *β*_1_ was set to 1 without loss of generality. However, in our aggregated neural activity model (3), when both the spiking rate *r* (given by the recorded spike count) and fluorescence *y* are provided to us, the relationship between them does not necessarily adhere to this assumption. To estimate *β*_1_ and express it explicitly in model (3), we can assume that the scaling it represents, between the calcium units and fluorescence unit, is the same for all neurons. Hence, expressing *β*_1_ explicitly in model (3) reads:

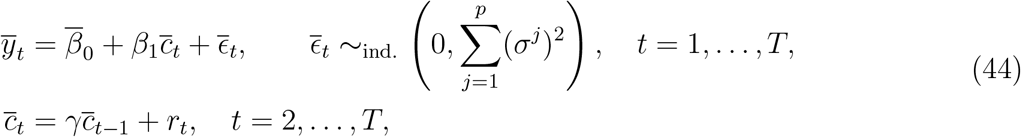

where 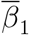 is the scaling between the calcium units and the fluorescence units. All other parameters are defined in Section 3.2. We can express the model above, excluding the noise contribution, using *A* and 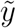 (defined in Section 3.3)

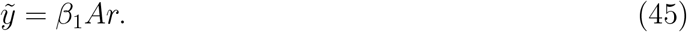

*β*_1_ is then the solution to the problem

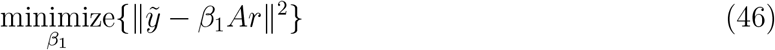

yielding

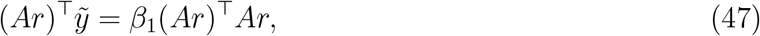

and the expression for *β*_1_

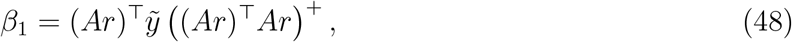

where ((*Ar*)^⊤^*Ar*) ^+^ is the pseudo inverse of (*Ar*)^⊤^*Ar*.

We recovered *β*_1_ for each 20sec segment in a dataset using (48). Segments with outlier values of *β*_1_ were excluded from further analysis (we visually verified that these segments were associated with technical errors in the recordings - evident in fluorescence traces that showed significantly increased magnitude or noise - or they represented periods of no correlation, or anti-correlation, between spike counts and fluorescence). We then scaled the recorded fluorescence *y* by 1*/ < β*_1_ *>* with *< β*_1_ *>* the mean value of *β*_1_ from all non-outlier segments in the dataset. This scaling ensured that *β*_1_ = 1 between the scaled fluorescence and spike counts. We used the scaled fluorescence (in the non outliers segments) as input to the deconvolution methods.

## S4 Calculating the Error and Choosing Penalty *λ* from Solely Fluorescence Recordings

Adopted from Jewell et al. (2019), Algorithm 3.

### Algorithm 3

A cross-validation scheme for choosing *λ*

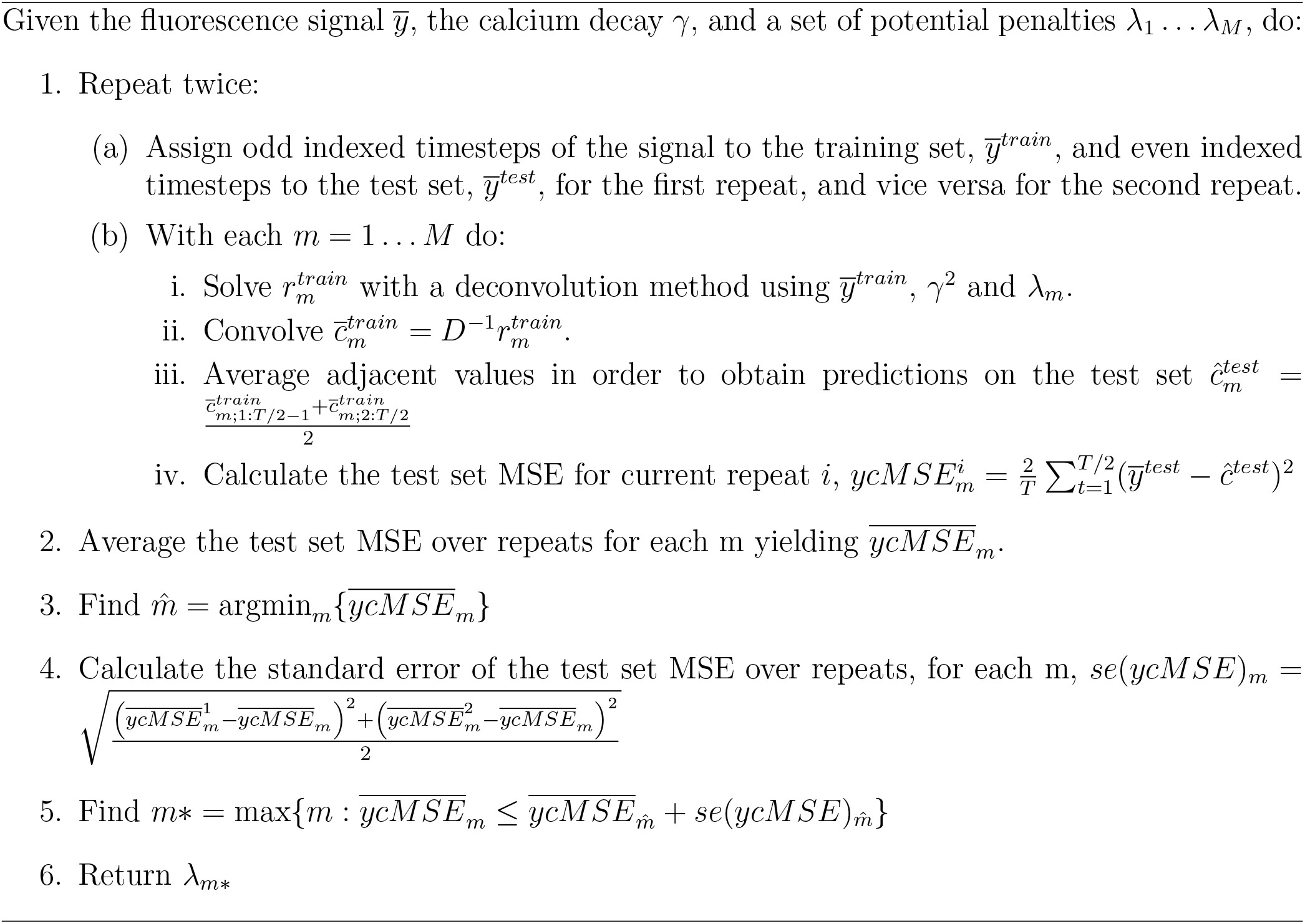

## Footnotes

1 To our knowledge, no other deconvolution method was adapted or developed for inferring aggregated neural activity captured by mesoscale recordings such as wide-field imaging. Other ad hoc approaches have been used, but their performances are significantly poorer, as expected.

2 An analytic solution was found in a parallel study (Stern 2024). Although the iterations of the algorithm proposed here require longer run times, the solutions are otherwise equivalent.

3 In a parallel work, an analytic solution to Continuously-Varying was found (Stern 2024). While the algorithm developed here converges to the same solution, using the analytic solution allows for a fast implementation of this method.

